# Diversity and connectivity of principal neurons in the lateral and basal nuclei of the mouse amygdala

**DOI:** 10.1101/2023.12.29.573684

**Authors:** Zsófia Reéb, Dániel Magyar, Filippo Weisz, Zsuzsanna Fekete, Kinga Müller, Attila Vikór, Zoltán Péterfi, Tibor Andrási, Judit M. Veres, Norbert Hájos

## Abstract

The basolateral amygdala is a non-layered cortical structure playing a role in various cognitive processes. Despite many studies focusing on local information processing within the circuits of the basolateral amygdala, the characteristics of excitatory principal neurons (PNs) are still not fully revealed. Here, we combined neuroanatomical, electrophysiological, and tracing techniques to determine the single-cell features, dendritic and axonal projections of PNs within the lateral (LA) and basal amygdala (BA). Using a mouse reporter line, we found that cholecystokinin (CCK) expression defines two spatially and functionally segregated groups of PNs both in the LA and BA. PNs in CCK-positive (CCK+) areas of the LA (LAa) had small somata and short dendrites which matched their single-cell electrophysiological properties. PNs in CCK-negative (CCK-) areas of the LA (LAp) and all BA had similarly ramified dendrites and single-cell features with some differences. Importantly, the dendritic arbors of PNs were restricted to the subnuclei defined by CCK expression, which corresponded to various extra-amygdalar afferents indicating specific inputs on distinct PN groups. Axonal arborization patterns of PNs within the basolateral amygdala and surrounding areas showed consistency to their soma location. For instance, BA PNs that projected to the medial prefrontal cortex but not to the lateral nucleus of the central amygdala were present in CCK+ areas. In contrast, those BA PNs that projected to the lateral part of the central nucleus were found in the subnucleus lacking CCK. Our study revealed that the basolateral amygdala is composed of functionally different subnuclei with specific inputs and outputs. This structural arrangement may empower the LA and BA to flexibly channel processed information toward their downstream regions, which can be a key requirement for diverse amygdala functions in cognitive operation.

## INTRODUCTION

The amygdala is a complex structure in the brain comprised of multiple interconnected nuclei playing roles in a broad spectrum of normal behavioral functions and pathological conditions (Janak & Tye, 2015; LeDoux, 2007). The basolateral amygdala complex (BLA) is a well-defined region, composed of the lateral (LA), basal (BA) and basomedial (BMA) nuclei (Pitkanen et al., 1997; Sah et al., 2003). The principal neurons (PNs) in the BLA are glutamatergic projecting cells (McDonald, 1982, 1992; McDonald & Culberson, 1981; Sah et al., 2003) accounting for ∼80% of all neurons in the mouse LA and BA (Vereczki et al., 2021) and sending axonal projections toward cortical and subcortical structures (Hintiryan et al., 2021; McDonald, 1992, 1998; Pitkanen, 2000; Sah et al., 2003). Although the BLA has a cortical origin based on the developmental studies (Puelles et al., 2000; Swanson & Petrovich, 1998), BLA PNs are not organized into layers (McDonald, 1982; Sah et al., 2003), posing a challenge in exploring their network organization principles.

BLA nuclei are heterogeneous in their morphological and electrophysiological characteristics or functionality (Pitkanen & Amaral, 1998; Pitkanen et al., 1997; Sah et al., 2003). For instance, the BA has been parceled by early neuroanatomical studies into anterior and posterior parts (McDonald, 1984; Pitkanen et al., 1995), which were supported by transcriptomic investigations (Kim et al., 2016). The most recent mapping of the brain-wide connectivity of the BA has further delineated the anterior BA to three domains: medial, lateral and caudal parts (Hintiryan et al., 2021). These and additional studies have highlighted that PNs with different projection areas spatially segregate within the BA (Beyeler et al., 2018; Hintiryan et al., 2021; McGarry & Carter, 2017; O’Leary et al., 2020). For instance, neurons targeting the prelimbic cortex (PL) involved in controlling fear states (Janak & Tye, 2015; Klavir et al., 2017; Senn et al., 2014) and anxiety (Felix-Ortiz et al., 2016; Liu et al., 2020; Lowery-Gionta et al., 2018) are more abundant in the anteromedial BA (Beyeler et al., 2018; Manoocheri & Carter, 2022; O’Leary et al., 2020), while central amygdala (CEA) projecting neurons conveying information of conditioned fear (Kim et al., 2017; Massi et al., 2023; Tye et al., 2011) are mainly located in the posterolateral BA (Beyeler et al., 2018; Massi et al., 2023). Interestingly, the mouse LA has not been divided further in these recent studies (Hintiryan et al., 2021), although previous studies in rats and monkeys proposed the presence of distinct subnuclei in the LA, too (Pitkanen, 2000; Pitkanen & Amaral, 1998; Pitkanen et al., 1997; Pitkanen et al., 1995; Sah et al., 2003).

Cholecystokinin (CCK) is a neuropeptide that has been shown to influence the local function of the BLA through altering fear-and anxiety-related behavior (Bowers & Ressler, 2015; Erlich et al., 2012; Truitt et al., 2009). In BAC-CCK-DsRed transgenic mice, DsRed fluorescent protein is expressed under the CCK promoter, resulting in fluorescent labeling of CCK containing neurons (Mate et al., 2013). Both PNs and a group of GABAergic basket cells express CCK in the BLA (Vereczki et al., 2016; Veres et al., 2017), with the former group characterized by weak and the latter by strong DsRed expression (Rovira-Esteban et al., 2017). The CCK promoter-driven expression of DsRed signals in PNs shows heterogeneity along the anteroposterior axis both within LA and BA, suggesting the presence of distinct domains in these BLA nuclei.

In the present study, first we analyzed the location of dendritic trees of *in vitro* filled neurons as input sites of the PNs, along with the pattern of some afferents innervating the LA and BA. Next, we explored the distribution of spatially segregated neuron populations within the LA and BA projecting to the prelimbic cortex (PL), dorsomedial striatum (DMS), central amygdala (CEA) and ventral hippocampus/subiculum (vHipp/SUB) in relation to the CCK-positive (CCK+) and CCK-negative (CCK-) areas. Finally, we performed the morphological and electrophysiological characterization of PNs of different LA and BA regions using *in vitro* and *in vivo* techniques. Our data demonstrate that CCK expression-defined areas within the LA and BA are functional subnuclei. In addition, we observed that PNs in CCK+ LA (i.e., in anterior LA) are smaller and have shorter dendritic arbors compared to those PNs located in the posterior LA and BA. Finally, our results revealed that the axons of PNs arborize stereotypically in the amygdala and the surrounding areas, identifying e.g., axonal collaterals of BA PNs in the LA.

## RESULTS

### CCK promoter-driven expression of DsRed defines LA and BA subnuclei

To study the distribution of CCK-expressing PNs in the BLA, BAC-CCK-DsRed transgenic mice were used (Mate et al., 2013; Rovira-Esteban et al., 2017). In low magnification images of coronal and horizontal amygdala sections obtained from these transgenic mice, distinct subunits can be separated both in the LA and BA based on the DsRed content of the neurons: one region shows strong DsRed signaling, i.e., CCK expression (CCK+ area), while the other part contains only sparsely labeled DsRed (i.e. CCK)-expressing neurons (CCK-area) (**Fig. 1A, B**). The CCK+ subnucleus is located in the anterior and lateral segments of the LA (LAa) and in the anterior and medial segments of the BA (BAa). Conversely, the posterior parts (LAp and BAp) are CCK-. Thus, DsRed/CCK expression in this transgenic mouse line clearly divides both the LA and the BA to two subnuclei.

**Fig. 1.**
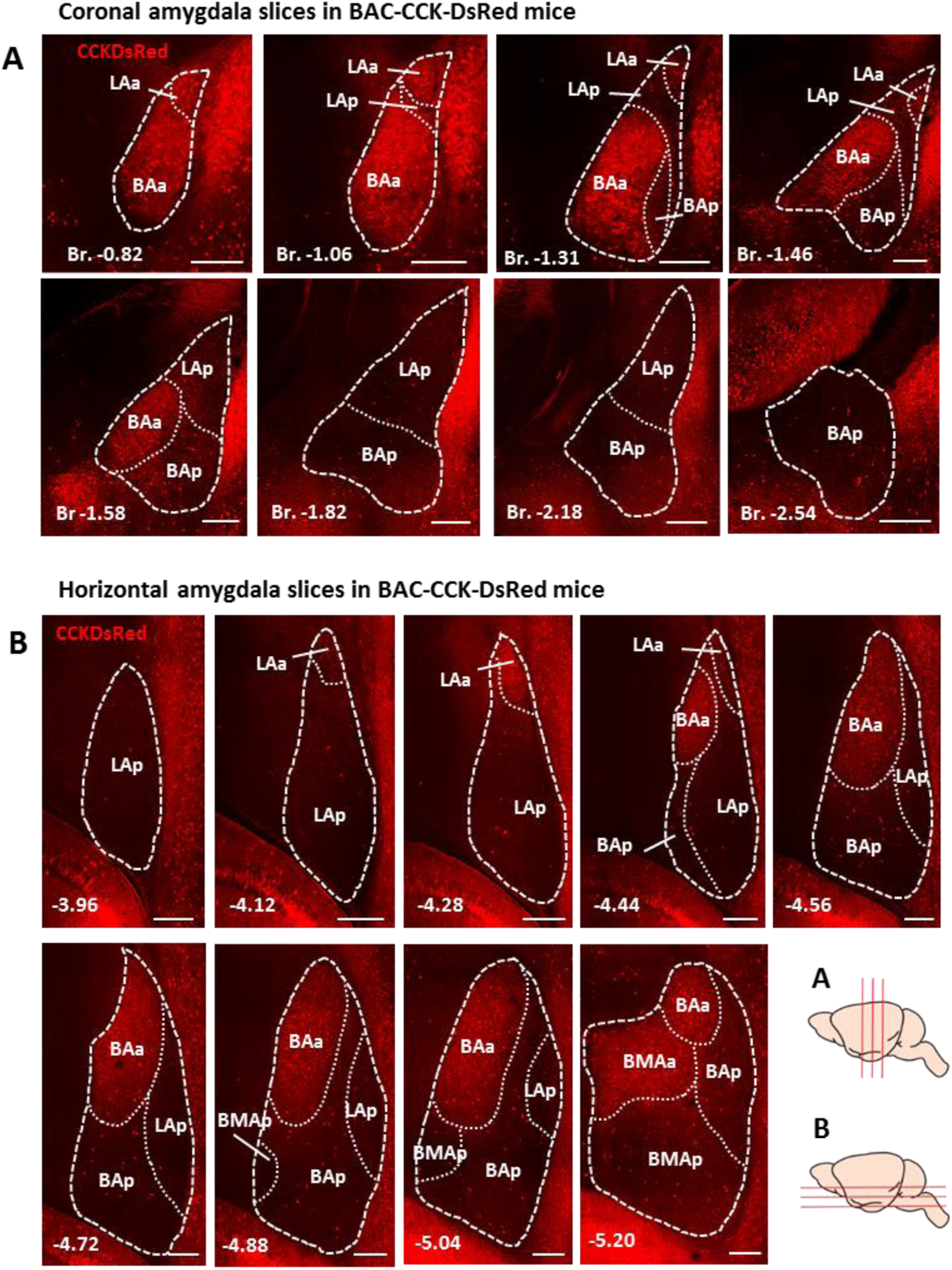
Cholecystokinin (CCK) promoter-driven expression of DsRed defines LA and BA subnuclei. (**A-B)** Confocal images of (**A)** coronal and (**B)** horizontal sections of the amygdala from BAC-CCK-DsRed mice indicate the distribution of endogenously expressing DsRed fluorescent protein under the control of CCK promoter. The density of CCK-expressing neurons clearly divides the LA and BA into a CCK+ anterior (LAa and BAa) and a CCK-posterior (LAp and BAp) part. White dashed and dotted lines represent the borders of the BLA and their subnuclei, respectively. Scale bar: 250 µm. Abbreviations: LAa: lateral amygdalar nucleus, anterior part; LAp: lateral amygdalar nucleus, posterior part; BAa: basal amygdalar nucleus, anterior part; BAp: basal amygdalar nucleus, posterior part; BMa: basomedial amygdalar nucleus, anterior part; BMp: basomedial amygdalar nucleus, posterior part; Br.: Bregma. In coronal slices numbers refer to the distance from Bregma in mm; in horizontal slices the interaural distance is indicated.

### The dendritic trees of PNs are restricted to the subnucleus where their cell bodies are located

To determine whether CCK expression-defined subnuclei are independent information processing units, we examined the location and extension of dendritic arborizations of PNs in coronal and horizontal slice preparations (**Fig. 2A**). PNs in different subnuclei were filled with biocytin followed by visualization of their biocytin content. After immunolabeling, the dendritic trees of individual PNs were manually reconstructed based on high-resolution confocal images and the proportion of dendritic arbors located within and outside of the CCK+ regions were determined. The borders between the LA and BA were determined based on the immunostaining against vesicular acetylcholine transporter (VAChT) (Arvidsson et al., 1997; Vereczki et al., 2021) (**Fig. S1)**. Our results show that the dendritic trees of the reconstructed PNs were restricted to the subnucleus of soma location (71-99% of the dendritic length was present in the given subnucleus on average). Accordingly, in the CCK+ LAa and BAa subnuclei, dendritic trees were confined to the areas defined by CCK expression, while in the CCK-LAp and BAp, they clearly avoided the CCK+ regions (**Fig. 2B-F**). These results imply that CCK expression can define functional units within the LA and BA, where the dendrites of PNs are limited to the given subnucleus.

**Fig 2.**
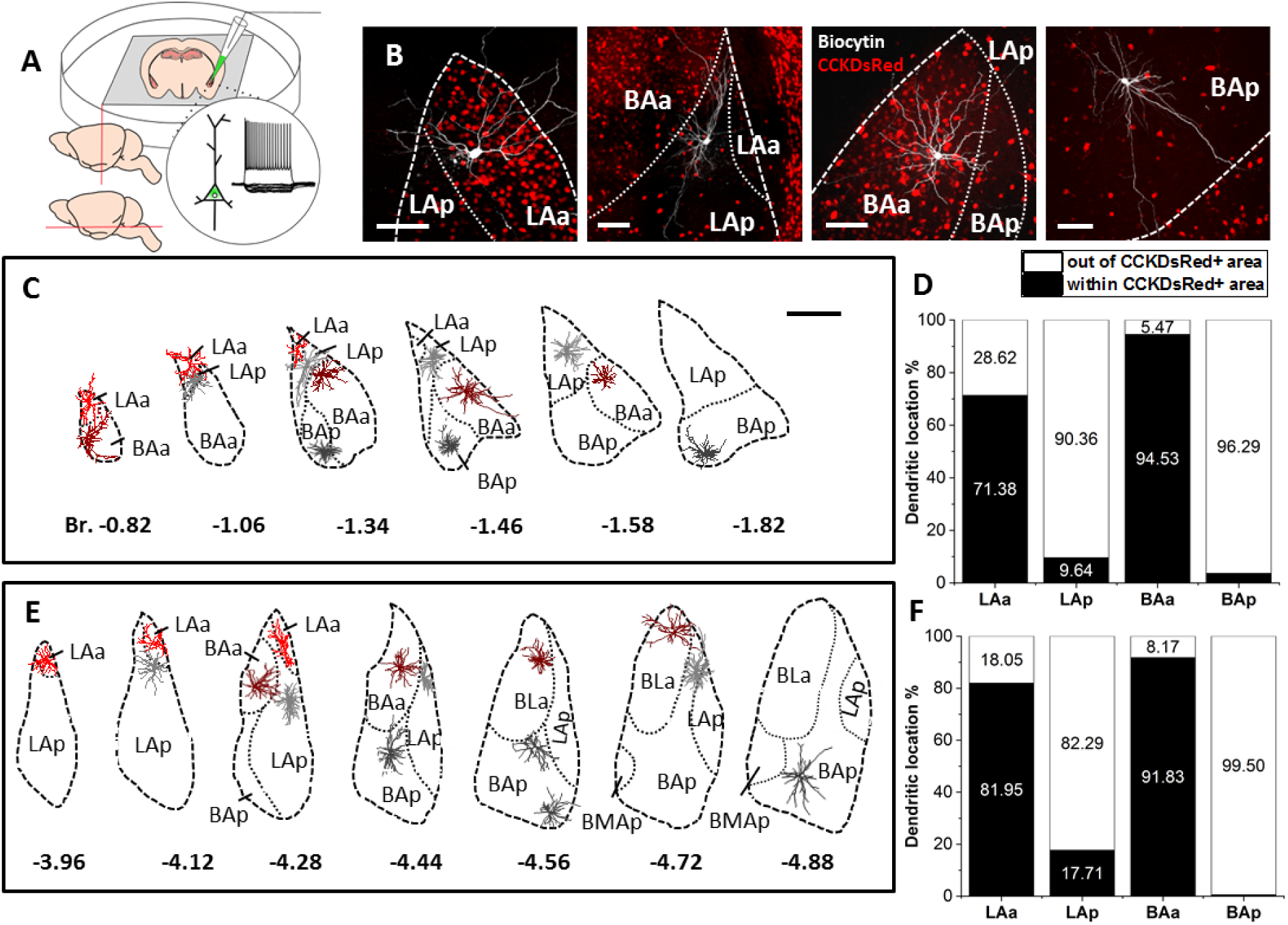
The dendritic trees of principal neurons show spatial restriction to the subnucleus where the soma is located. (**A)** Experimental setup. *In vitro* whole-cell recording and labeling of PNs in coronal and horizontal slice preparations, focusing on distinct subnuclei of LA and BA. (**B)** Confocal images of biocytin filled example neurons showing restricted dendritic trees to LA and BA subnuclei. Images of example PNs taken from coronal (LAp PN and BAp PN) and horizontal (LAa PN and BAa PN) slices. Scale bar: 100 µm. (**C)** Reconstructed dendritic trees of 14 representative PNs from LA and BA subnuclei obtained from 6 different coronal sections along the antero-posterior axis of the amygdala. Scale bar: 500 µm. (**D)** Bar graph showing the proportion of dendritic trees located within and outside of CCK+ areas in coronal slices (LAa: n=9 (red); LAp: n=8 (light gray); BAa: n=9 (dark red) and BAp: n=9 (dark gray); 35 PNs from 17 mice). Dendritic trees of PNs with somata located in the LAa and BAa are restricted to CCK+ areas. Conversely, dendritic trees of PNs with somata located in the LA or BAp are restricted to CCK-areas. (**E)** Reconstructed dendritic trees of 15 representative PNs from the LA and BA subnuclei obtained from 7 different horizontal sections along the dorso-ventral axis of the amygdala. (**F)** Bar graph showing the proportion of dendritic trees located within and outside of the CCK+ areas in horizontal slices (LAa: n=8 (red); LA: n=13 (light gray); BAa: n=9 (dark red) and BAp: n=23 (dark gray); 53 PNs from 23 mice). Dendritic trees of PNs with somata located in the LAa and BAa are restricted to CCK+ areas. Conversely, dendritic trees of PNs with somata located in the LA or BAp are restricted to CCK-areas. Scale bar: 500 µm.

### CCK expression-defined subnuclei can be considered as functional units

If the subregions defined by CCK expression form functional subunits, then it is likely that these distinct subnuclei receive different sets of afferents that specify the inputs received by PNs with subregion-confined dendritic trees. To test this hypothesis, we examined whether the termination pattern of afferents reaching the amygdala follows the borders of CCK expression-defined subnuclei. The medial prefrontal cortical (mPFC), the insular cortical (IC) and the midline thalamic (MT) afferents were investigated, as these are among the main sources of inputs to the LA and BA (**Fig. 3**., **S2-S4**). AAV9-Ef1a-DIO-eYFP was injected into the mPFC or IC of Vglut1-Cre::BAC-CCK-DsRed mice expressing Cre recombinase under the promoter of vesicular-glutamate transporter type 1 (vGlut1) and DsRed fluorescent protein under the CCK promoter (n=2-3). This approach allowed us to trace the cortical glutamatergic inputs reaching the amygdala and compare their pattern to CCK expression. In rostral and caudal segments of the basolateral amygdala both mPFC and IC afferents targeted the BAa and the BAp but in a complementary manner (**Fig. S2, S3**). In the medial segments, the BAa was exclusively innervated by the mPFC afferents, whereas the LAp and BAp subnuclei were innervated by the IC axons (**Fig. 3A-B**, **S2, S3**). Since there is no vGlut1 expression in the thalamus (Fremeau et al., 2001), MT afferents were labeled using AAV5-CaMKII-eYFP (n=2). The axons of these thalamic inputs primarily targeted the BA, regardless of the subunits determined by CCK expression (**Fig. 3**, **S3**) in line with previous observations (Ahmed et al., 2021; Amir et al., 2019; Matyas et al., 2018). These results demonstrate that CCK expression in PNs defines functional units in the LA and BA as both the dendritic trees and cortical inputs match the subnuclei delineated by DsRed signal expression in the PNs.

**Fig. 3.**
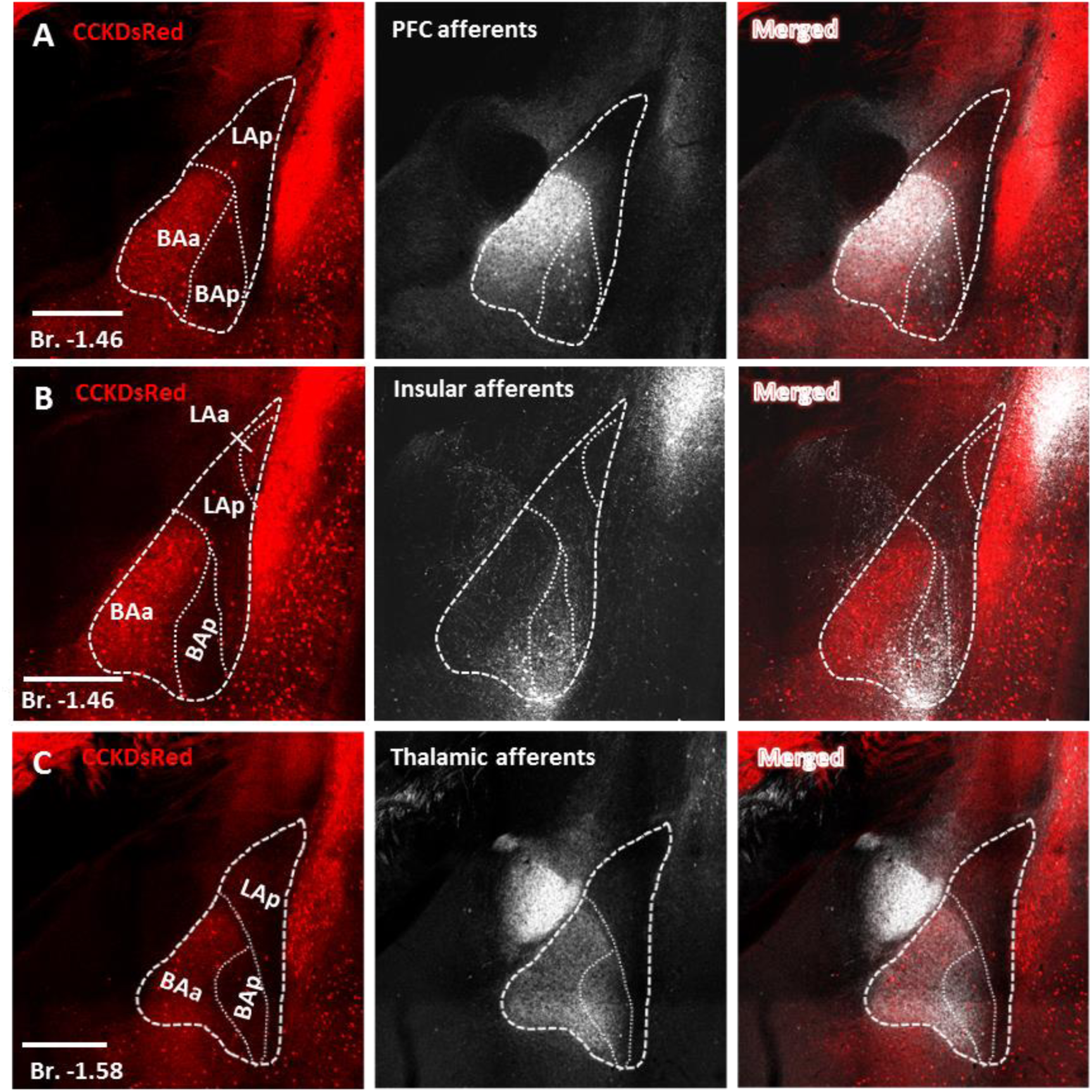
CCK-defined BA subnuclei receive distinct cortical inputs. (**A**) Prefrontal cortical (PFC), (**B**) insular cortical (IC), and (**C**) midline thalamic (MT) afferents innervating the amygdala in BAC-CCK-DsRed animals. PFC and IC target the BAa and BAp in a complementary manner, while axons of the MT primarily target both parts of the BA, regardless of the subunits determined by CCK expression. Scale bar: 500 µm. Numbers refer to the distance from Bregma (Br) in mm.

### Principal neurons of the LAa are smaller than those in the LAp

Next, we asked whether the PNs in distinct subnuclei in the LA and BA have different morphological and electrophysiological properties. To this end, we compared the features of *in vitro* filled PNs. (**Fig. 4**). LAa PNs were found to be smaller, having on average shorter dendritic length and less ramified dendrites than in other PNs examined in the CCK-part of the LA, in the LAp (**Fig. 4B, D**). Accordingly, significant differences were observed in the length of 4^th^ and 5^th^ order dendrites of PNs (**Fig. 4C**). Furthermore, Sholl analysis revealed that in 50-100 µm distance from the soma, the dendritic length of LAa PNs was shorter compared to PNs in other subnuclei (**Fig.4E**). As these PN labeling took place in slice preparations using whole-cell recordings allowing to study single-cell features, we could determine the active and passive membrane properties of PNs located in distinct subnuclei (**Table 1**). Positive and negative step currents were injected into the PN somata and the voltage responses were recorded and subsequently analyzed (**Fig. 5A**). In accord with the morphological data, LAa PNs had the lowest capacitance in comparison to other PNs (**Fig. 5B**). The input resistance of LAa PNs was found to be larger only in comparison to BAp PNs (**Fig. 5C**), whereas the membrane time constant, which determines the integration properties of neurons (Sprouston, 2016), was slower for LAp PNs in comparison to LAa and BAp PNs (**Fig. 5D**). Interestingly, the amplitude of the sag, which corresponds to the expression of h current (Maccaferri & McBain, 1996; Tanaka et al., 2003), was not different in PNs studied (**Fig. 5E**), showing that postsynaptic potentials may be controlled similarly along the dendrites by this active conductance (Magee, 1998). Lastly, we compared the spiking features and found that LAa PNs have the highest spike threshold in comparison to BA PNs (**Fig. 5F**) and the steepest input-output function (**Fig. 5G**). Thus, LAa PNs are overall the smallest PNs in the LA and BA and they can be activated by the smallest excitatory inputs. Dendritic morphology and single-cell features of BAa and BAp PNs did not differ.

**Fig. 4.**
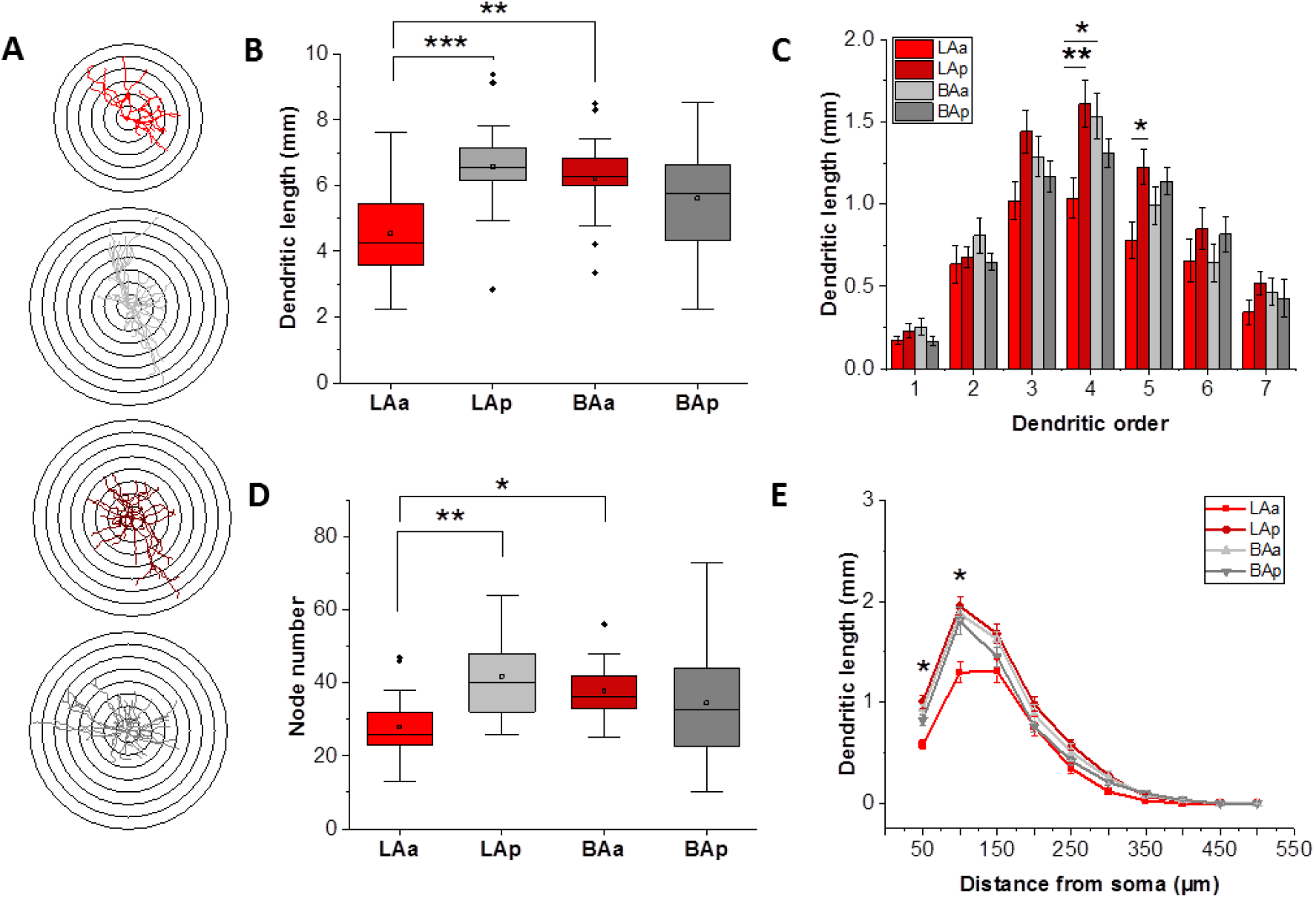
Dendritic tree features of principal neurons in the subnuclei of LA and BA. (**A**) Reconstructions of dendritic trees and Sholl-analysis of representative principal neurons (PNs) in each LA and BA subnucleus. The difference between the radii of concentric circles is 50 µm. (**B**) Comparison of total dendritic length from LA and BA subnuclei. (**C**) Comparison of the dendritic length of LAa, LA, BAa and BAp PNs as a function of dendritic order. Dunn’s test showed significant differences for 4^th^ (LAa–LAp: p=0.015; LAp–BAa: p=0.042) and 5^th^ (LAa–LAp: p=0.021) order dendrites. (**D**) Comparison of the number of branching nodes of dendritic trees of different PN groups. (**E**) Sholl analysis of reconstructed dendritic trees showing the dendritic length as a function of the distance from the soma. Dunn’s test revealed significant differences at 0-50 µm (LAa–LAp: p<0.001; LAa–BAa: p<0.001; LAa–BAp: p=0.021) and at 50-100 µm distance from the soma: LAa–LAp: p<0.001; LAa–BAa: p=0.007; LAa-BAp: p=0.016). (**A-E**) Data obtained in coronal and horizontal slices (LAa: n=17; LAp: n=21; BAa: n=18; BAp: n=32. 88 PNs from 40 mice). Statistical comparisons were performed with Kruskal-Wallis ANOVA followed by *post hoc* Dunn’s test. For detailed morphological parameters see **Table 2**.

**Fig. 5.**
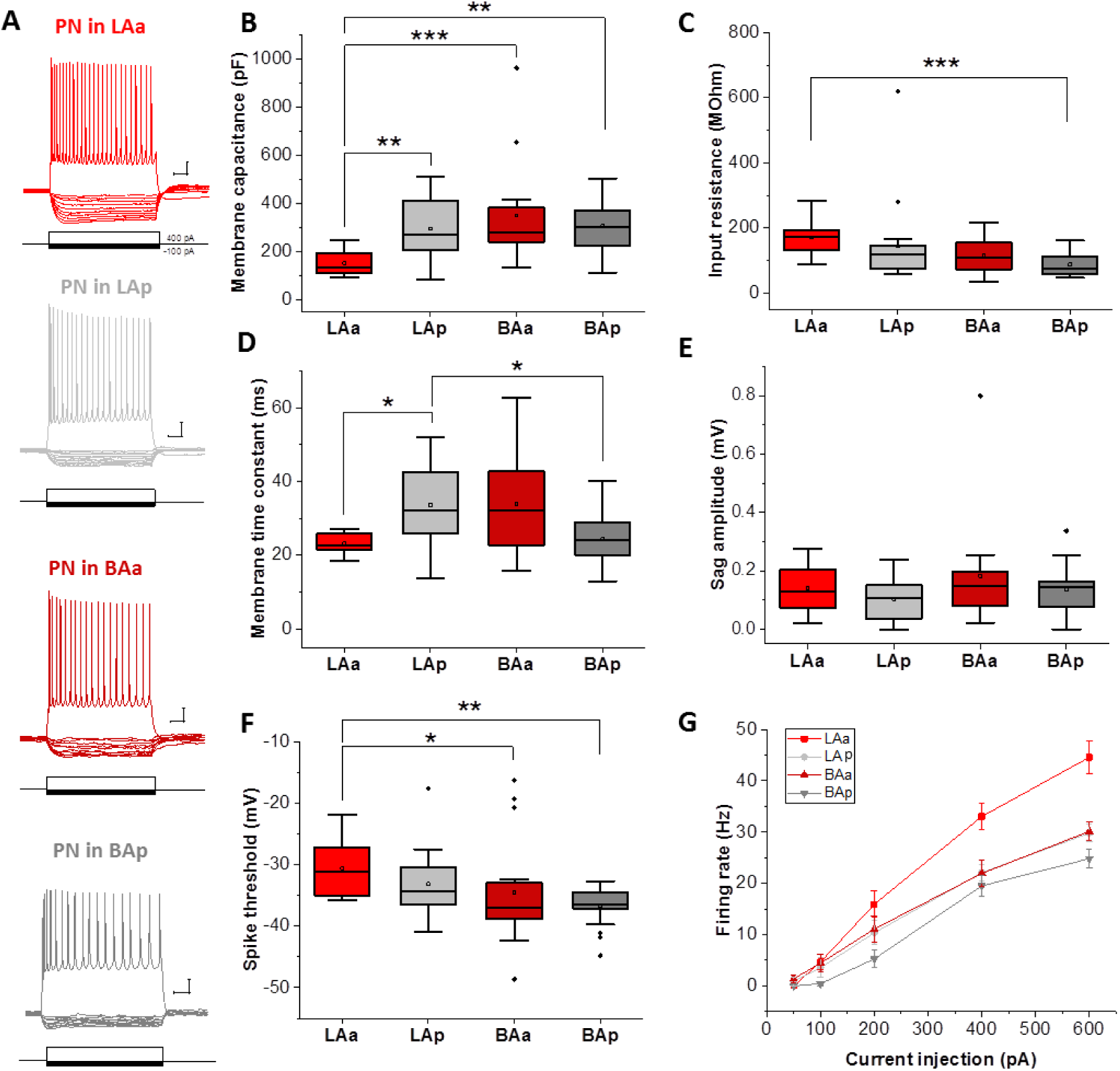
The active and passive membrane properties of *in vitro* recorded principal neurons in LA and BA subnuclei. (**A**) Example firing traces of principal neurons from each LA and BA subnuclei obtained in BA-CCK-DsRed animals. Scale bar: y: 10 mV and x: 100 ms. Box chart comparison of (**B**) membrane capacitance, (**C**) input resistance, (**D**) membrane time constant (**E**) sag amplitude and (**F**) spike threshold of the PNs in different LA and BA subnuclei (LAa: n=12; LAp: n=19; BAa: n=16; BAp: n=18-20. 67 PNs from 39 mice). Statistical comparisons were performed with Kruskal-Wallis ANOVA followed by *post hoc* Dunn’s test. (**G**) Firing rate as a function of injected current (LAa: n=6; LAp: n=3, BAa: n=6, BAp: n=15). Data are presented as mean ± SEM. See **Table 1** for further electrophysiological data and significance levels.

**Table 1.** Electrophysiological properties of principal neurons in different LA and BA subnuclei. Data are presented as the median with the first and third quartiles in parentheses. Significant differences shown in bold were determined by Kruskal-Wallis ANOVA and post hoc Dunn’s test (AP: action potential, AHP: after hyperpolarization).

|  | LAa<br>(n=10-12) | LAp<br>(n=15-19) | BAa<br>(n=12-16) | BAp<br>(n=17-20) | Kruskal-Wallis<br>ANOVA | LAa-LAp<br>Dunn's test | LAa-BAa<br>Dunn's test | LAa-BAp<br>Dunn's test | LAp-BAa<br>Dunn's test | LAp-BAp<br>Dunn's test | BAa-BAp<br>Dunn's test |
| --- | --- | --- | --- | --- | --- | --- | --- | --- | --- | --- | --- |
| **AP threshold**<br>**(mV)** | -31.10<br>(-35.10,<br>-26.40) | -34.30<br>(-36.40,<br>-30.40) | -37.00<br>(-39.12,<br>-32.62) | -36.40<br>(-37.10,<br>-34.50) | **0.003** | 0.842 | **0.021** | **0.005** | 0.551 | 0.193 | 1 |
| **Spike rate**<br>**(Hz)** | 38.80<br>(33.45,<br>53.12) | 31.30<br>(23.80,<br>36.30) | 30.65<br>(25.05,<br>36.30) | 25.65<br>(22.50,<br>28.80) | **<0.001** | 0.061 | 0.150 | **<0.001** | 1 | 0.623 | 0.404 |
| **Accommodation**<br>**ratio** | 3.91<br>(2.81,<br>5.25) | 4.96<br>(3.84,<br>5.65) | 6.49<br>(4.40,<br>7.55) | 5.21<br>(3.32,<br>6.83) | 0.056 |  |  |  |  |  |  |
| **Full width at half**<br>**maximum (ms)** | 0.65<br>(0.60,<br>0.90) | 0.60<br>(0.60,<br>0.80) | 0.70<br>(0.60,<br>0.87) | 0.90<br>(0.80,<br>0.90) | **0.007** | 1 | 1 | 0.178 | 1 | **0.005** | 0.329 |
| **AHP amplitude**<br>**(mV)** | 0.70<br>(0.60,<br>10.80) | 5.50<br>(0.90,<br>11.20) | 7.45<br>(3.67,<br>21.35) | 11.40<br>(8.67,<br>51.85) | **0.003** | 1 | 0.709 | **0.004** | 1 | **0.027** | 0.384 |
| **AHP 25% decay**<br>**(ms)** | 31.50<br>(2.40,<br>80.80) | 49.05<br>(7.57,<br>84.97) | 3.40<br>(1.70,<br>56.95) | 57.60<br>(40.20,<br>83.60) | 0.063 |  |  |  |  |  |  |
| **AHP 50% decay**<br>**(ms)** | 43.50<br>(4.60,<br>124.30) | 68.00<br>(14.55,<br>106.62) | 38.05<br>(3.77,<br>112.72) | 73.10<br>(58.40,<br>131.30) | 0.198 |  |  |  |  |  |  |
| **AHP 75% decay**<br>**(ms)** | 45.15<br>(26.37,<br>106.85) | 75.40<br>(31.00,<br>117.60) | 34.85<br>(16.57,<br>119.42) | 83.90<br>(62.90,<br>151.00) | 0.109 |  |  |  |  |  |  |
| **Sag amplitude** | 0.13<br>(0.07,<br>0.20) | 0.11<br>(3.21E-02, 0.16) | 0.15<br>(0.08,<br>0.20) | 0.14<br>(7.23E-02, 0.18) | 0.442 |  |  |  |  |  |  |
| **Input resistance**<br>**(MOhm)** | 173.00<br>(120.25,<br>197.15) | 118.50<br>(73.59,<br>144.20) | 107.75<br>(68.79,<br>159.40) | 76.15<br>(57.02,<br>159.40) | **<0.001** | 0.181 | 0.108 | **<0.001** | 1 | 0.156 | 0.417 |
| **Membrane**<br>**capacitance (pF)** | 130.80<br>(105.22,<br>205.13) | 267.20<br>(206.10,<br>409.20) | 278.90<br>(233.72,<br>398.60) | 300.00<br>(220.70,<br>375.48) | **<0.001** | **0.003** | **<0.001** | **0.001** | 1 | 1 | 1 |
| **Membrane time**<br>**constant (ms)** | 22.63<br>(21.34,<br>25.88) | 32.08<br>(25.96,<br>42.51) | 32.03<br>(22.58,<br>45.01) | 24.02<br>(20.06,<br>29.03) | **0.005** | **0.021** | 0.113 | 1 | 1 | **0.045** | 0.259 |

**Table 2.**
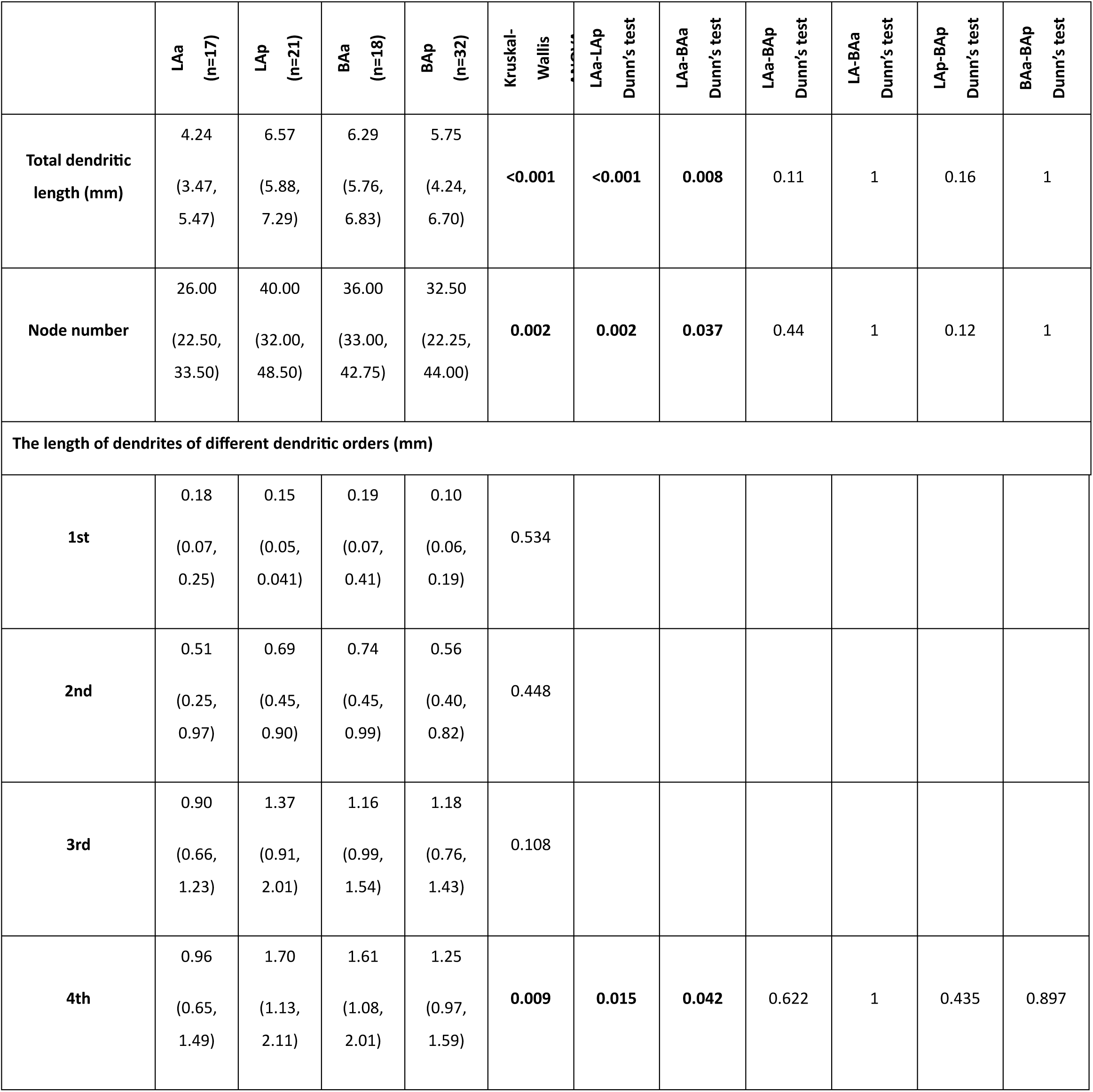
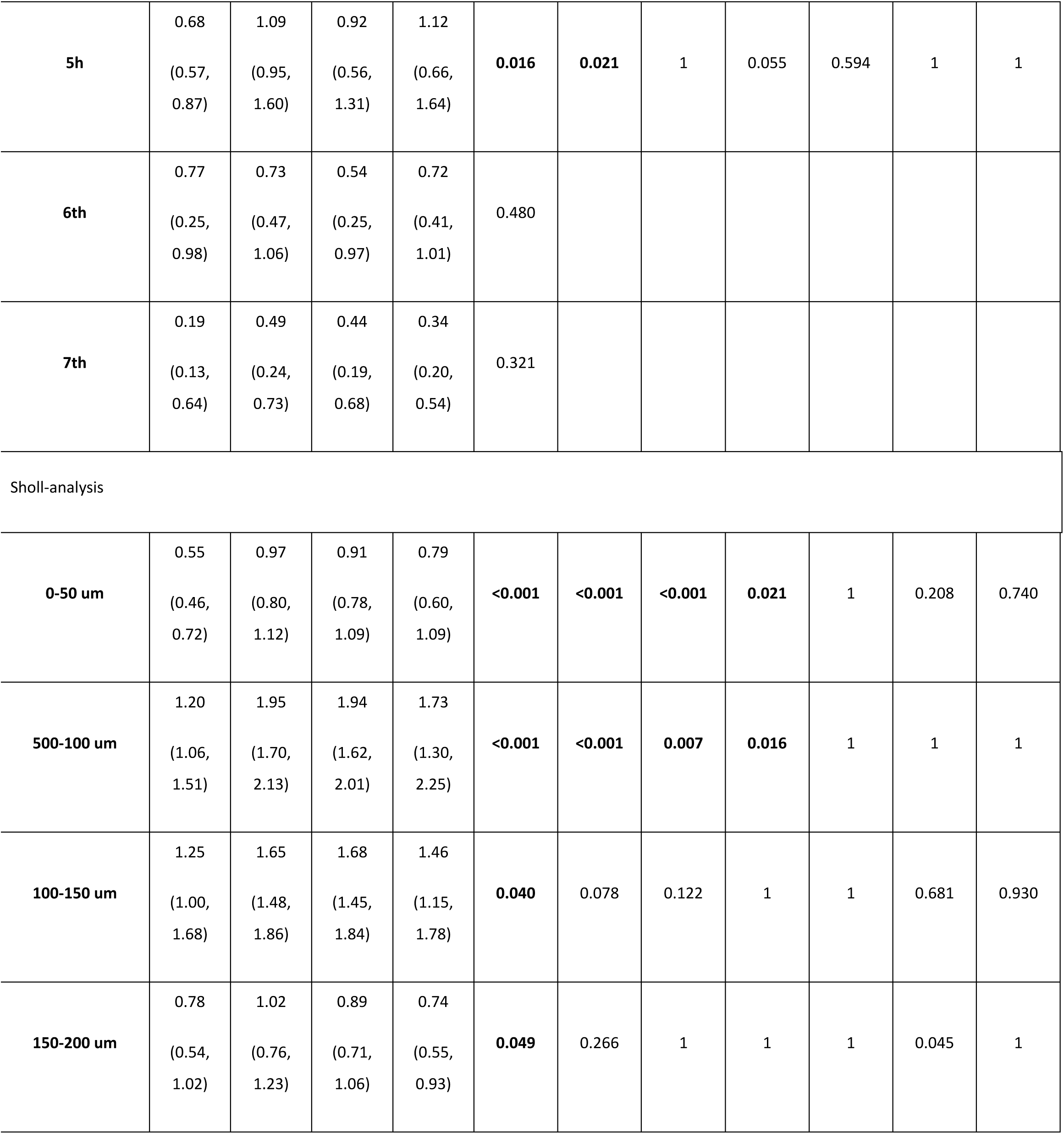
Morphological properties of the principal neurons in the different LA and BA subnuclei. Data are presented as the median with the first and third quartiles in parentheses. Significant differences shown in bold were determined by Kruskal-Wallis ANOVA and post hoc Dunn’s test.

|  | LAA<br>(n=17) | LAP<br>(n=21) | BAA<br>(n=18) | BAP<br>(n=32) | Kruskal-Wallis | LAA-LAP<br>Dunn's test | LAA-BAA<br>Dunn's test | LAA-BAP<br>Dunn's test | LA-BAA<br>Dunn's test | LAP-BAP<br>Dunn's test | BAA-BAP<br>Dunn's test |
| --- | --- | --- | --- | --- | --- | --- | --- | --- | --- | --- | --- |
| **Total dendritic length (mm)** | 4.24<br>(3.47, 5.47) | 6.57<br>(5.88, 7.29) | 6.29<br>(5.76, 6.83) | 5.75<br>(4.24, 6.70) | **<0.001** | **<0.001** | **0.008** | 0.11 | 1 | 0.16 | 1 |
| **Node number** | 26.00<br>(22.50, 33.50) | 40.00<br>(32.00, 48.50) | 36.00<br>(33.00, 42.75) | 32.50<br>(22.25, 44.00) | **0.002** | **0.002** | **0.037** | 0.44 | 1 | 0.12 | 1 |
| **The length of dendrites of different dendritic orders (mm)** |  |  |  |  |  |  |  |  |  |  |  |
| **1st** | 0.18<br>(0.07, 0.25) | 0.15<br>(0.05, 0.041) | 0.19<br>(0.07, 0.41) | 0.10<br>(0.06, 0.19) | 0.534 |  |  |  |  |  |  |
| **2nd** | 0.51<br>(0.25, 0.97) | 0.69<br>(0.45, 0.90) | 0.74<br>(0.45, 0.99) | 0.56<br>(0.40, 0.82) | 0.448 |  |  |  |  |  |  |
| **3rd** | 0.90<br>(0.66, 1.23) | 1.37<br>(0.91, 2.01) | 1.16<br>(0.99, 1.54) | 1.18<br>(0.76, 1.43) | 0.108 |  |  |  |  |  |  |
| **4th** | 0.96<br>(0.65, 1.49) | 1.70<br>(1.13, 2.11) | 1.61<br>(1.08, 2.01) | 1.25<br>(0.97, 1.59) | **0.009** | **0.015** | **0.042** | 0.622 | 1 | 0.435 | 0.897 |
| **5h** | 0.68<br>(0.57,<br>0.87) | 1.09<br>(0.95,<br>1.60) | 0.92<br>(0.56,<br>1.31) | 1.12<br>(0.66,<br>1.64) | **0.016** | **0.021** | 1 | 0.055 | 0.594 | 1 | 1 |
| **6th** | 0.77<br>(0.25,<br>0.98) | 0.73<br>(0.47,<br>1.06) | 0.54<br>(0.25,<br>0.97) | 0.72<br>(0.41,<br>1.01) | 0.480 |  |  |  |  |  |  |
| **7th** | 0.19<br>(0.13,<br>0.64) | 0.49<br>(0.24,<br>0.73) | 0.44<br>(0.19,<br>0.68) | 0.34<br>(0.20,<br>0.54) | 0.321 |  |  |  |  |  |  |
| Sholl-analysis |  |  |  |  |  |  |  |  |  |  |  |
| **0-50 um** | 0.55<br>(0.46,<br>0.72) | 0.97<br>(0.80,<br>1.12) | 0.91<br>(0.78,<br>1.09) | 0.79<br>(0.60,<br>1.09) | **<0.001** | **<0.001** | **<0.001** | **0.021** | 1 | 0.208 | 0.740 |
| **500-100 um** | 1.20<br>(1.06,<br>1.51) | 1.95<br>(1.70,<br>2.13) | 1.94<br>(1.62,<br>2.01) | 1.73<br>(1.30,<br>2.25) | **<0.001** | **<0.001** | **0.007** | **0.016** | 1 | 1 | 1 |
| **100-150 um** | 1.25<br>(1.00,<br>1.68) | 1.65<br>(1.48,<br>1.86) | 1.68<br>(1.45,<br>1.84) | 1.46<br>(1.15,<br>1.78) | **0.040** | 0.078 | 0.122 | 1 | 1 | 0.681 | 0.930 |
| **150-200 um** | 0.78<br>(0.54,<br>1.02) | 1.02<br>(0.76,<br>1.23) | 0.89<br>(0.71,<br>1.06) | 0.74<br>(0.55,<br>0.93) | **0.049** | 0.266 | 1 | 1 | 1 | 0.045 | 1 |

### The distribution of principal neuron somata with different projection sites within the LA and BA corresponds to the CCK expression-defined functional subunits

Several studies have highlighted that BLA PNs with different projection targets are spatially segregated (Beyeler et al., 2018; Hintiryan et al., 2021; McGarry & Carter, 2017; O’Leary et al., 2020). Therefore, we examined how the distribution of these neuronal populations relates to the subnuclei defined by CCK expression. We addressed this question by retrograde labeling of those neurons that project to the distinct target areas of the basolateral amygdala. Cholera toxin B subunit (CTB), a retrograde tracer or retrograde adeno-associated virus (AAVrg) was injected into the PL, DMS, CEA and vHipp/SUB, respectively (**Fig. S5**), to reveal the PN populations with different projection targets in the LA and BA. To achieve more complete labeling, tracers were administered at two anteroposterior coordinates (except for the CEA due to its small size). Our data demonstrate that PL-and DMS-projecting PNs were located almost exclusively in the BAa (**Fig. 6A**), while the somata of CEA-projecting neurons were restricted to the BAp in addition to the LA (**Fig. 6B**). PNs projecting to the vHipp/SUB did not show spatial segregation within the BA (**Fig. 6C**). Similarly, spatial segregation within the LA was not observed in case of either projections (**Fig. 6A-C**). Finally, we examined to what extent the PN populations projecting to the PL and DMS overlap within the BAa. To this end, two different rgAAVs expressing distinct reporter proteins were injected to the PL and DMS, respectively. In addition to the similar ratio of single labeled neurons (PL-projecting PNs: 40.8%, 269/659; DMS-projecting PNs: 45.5%, 300/659, n=2 mice), there were numerous co-labeled PNs in the BAa (13.7%, 90/659), suggesting the presence of a PN population that sends collaterals to both brain regions (**Fig. 6A**). To confirm the collateralization and extend these findings, an intersectional approach was used to examine the axonal projections of PNs that innervated the PL and DMS (**Fig. S6A, E**). AAVrg-Cre was injected to the PL and DMS, respectively, whereas AAV-DIO-EYFP was injected to the BA. This approach allowed labeling the BA PNs projecting to the PL or in separate experiments to the DMS and their axonal collaterals. We observed axons of PL-projecting BA PNs in the DMS, too, however, no collateral was found in the lateral nucleus of the CEA (**Fig. S6B-D**). Similarly, axon collaterals of DMS-projecting BA PNs were also found in the PL and the anterior cingulate cortex, primarily in layers II/III and Vb; whereas, no fibers were revealed in the lateral nucleus of the CEA (**Fig. S6G-J**). These findings confirm the results obtained by double retrograde tracing, indicating that there is a population of BA PNs that project both to the PL and DMS, and suggest that the axons of both PL-and DMS-projecting BA PNs avoid the lateral nucleus of the CEA.

**Fig. 6.**
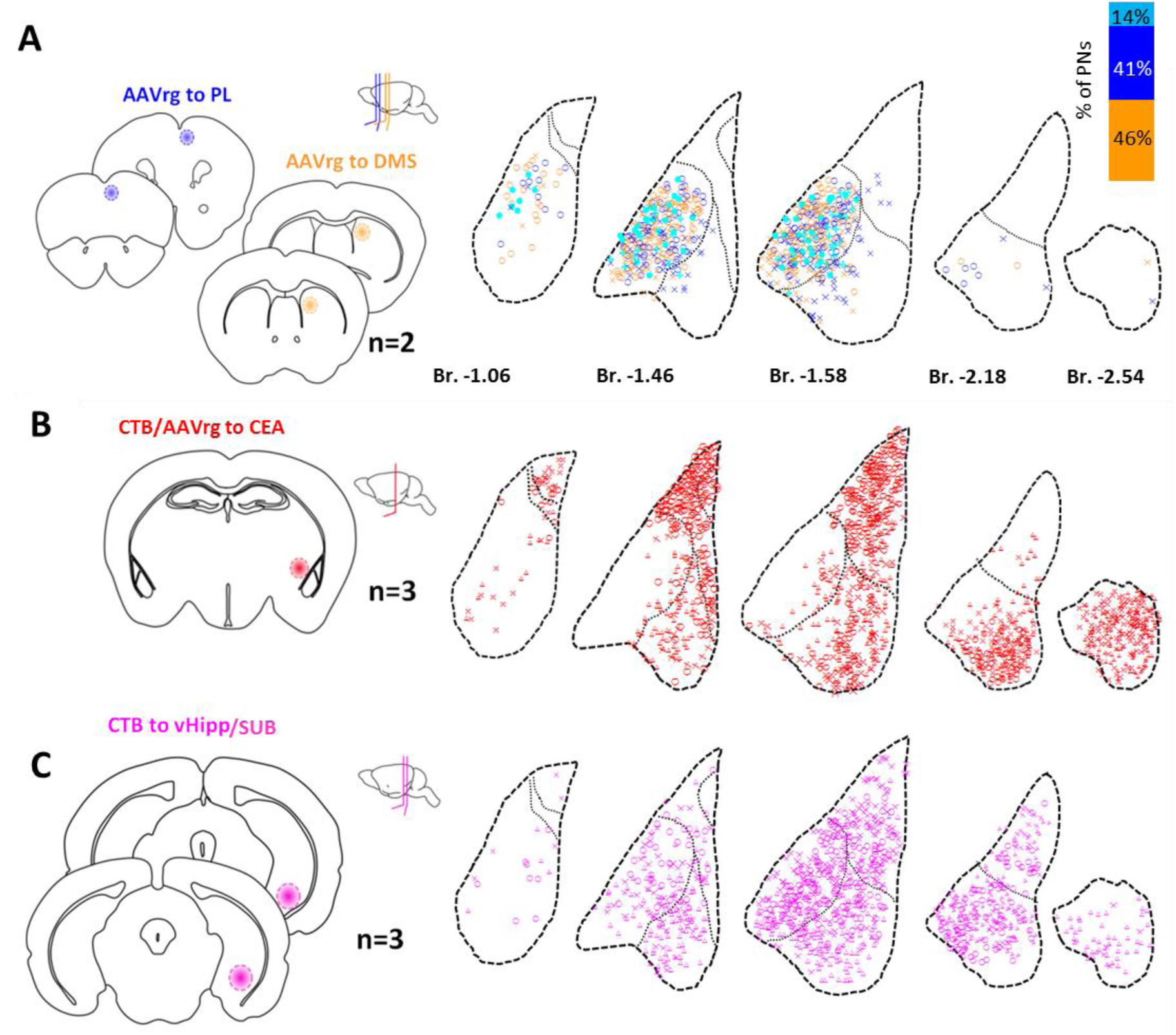
The distribution of the somata of principal neurons with different projection sites within the LA and BA corresponds to the CCK-DsRed defined functional subnuclei. (**A**) Left panels show the schematic representation of injection sites in the prelimbic cortex (PL) and dorsomedial striatum (DMS) targeted with two different types of retroAAV (AAVrg), respectively. The insert displays the sections of the mouse brain containing the core of the injection sites (2 injections/brain area). On the right, retrogradely labeled neurons are presented in five LA and BA sections (n=2 mice). PL-projecting neurons are shown in dark blue, while DMS-projecting in orange. Cyan stands for the co-labeled neurons sending collaterals to both brain regions. The bar graph in the right upper corner shows the ratio of PL-, DMS-and dual-projecting PNs, counted in 2 animals. (**B-C**) Left panels show the schematic representations of Cholera toxin B subunit (CTB)/retrograde AAV injection sites to the (**B**) central amygdala (CEA) or (**C**) ventral hippocampus (vHipp)/subiculum (SUB). Left schematics indicate the injection sites, while right panels show the retrogradely labeled neurons in five sections of the LA and BA (n=3 mice/target site). Thinner black dashed lines show the borders of the LA and BA subnuclei, different markers (circle, triangle, X) represent the results from 3 different animals. The distance from Bregma (Br.) is shown in mm.

Overall, our results suggest that the CCK expression segregates the areas where PL-and/or DMS-projecting PNs and CEA-projecting PNs are found in the BA, whereas BA PNs that project to the vHipp/SUB are present in both subnuclei. In addition, there is no spatial segregation of PNs in the LA that project to the CEA or vHipp/SUB.

### Reconstructions of *in vivo* filled PNs provide unique examples of connections between neighboring brain areas

Tracing studies are well-suited for revealing the long-range projections of PNs, but this method is only partially suited to reveal their local axonal arborizations. Knowing the local connectivity, however, is essential to understand the logic of intra-amygdalar information processing. Therefore, we performed *in vivo* juxtacellular recordings in the LA and BA in anesthetized mice (**Fig. 7A**), a single-cell labeling technique that allows visualizing the entire dendritic arbors and axonal projections of individual neurons (Pinault, 1996; Pitkanen et al., 2003). From 100 labeled neurons (n=55 mice), 21 were chosen for reconstructions that were identified as PNs with well-labeled spiny dendritic trees and axons that showed no or minimal weakening in staining intensity even at the distal parts of their arborizations. First, we examined the morphological properties of the dendritic trees of *in vivo* filled PNs (**Fig. 7**). By comparing the total dendritic length, structure, and the number of branching points of 21 PNs from different subunits, we observed that PNs in the LAp (n=4), BAa (n=3) and BAp (n=12) have similar dendritic arborizations (**Fig. 7B-E**). In addition, we successfully labeled 2 PNs in the LAa that had shorter dendritic length and correspondingly less branch points (**Fig. 7B-E**). Overall, these data obtained *in vivo* are in a good agreement with those obtained in slice preparations, indicating that LAa PNs are smaller with less elaborated dendritic trees in comparison to PNs in the LAp, BAa and BAp, which have similar dendritic morphology.

**Fig. 7.**
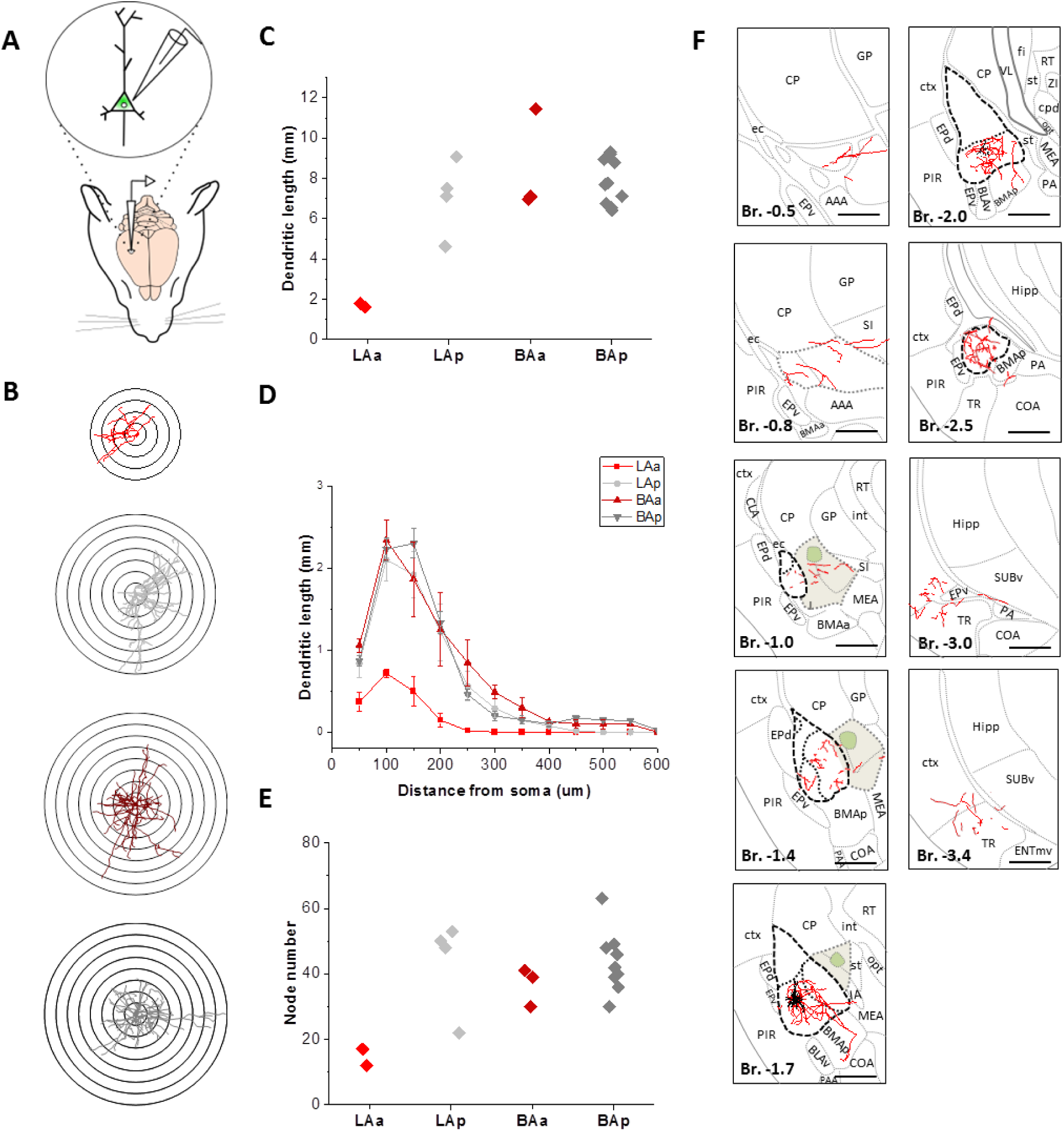
*In vivo* juxtacellular labeling of principal neurons in the LA and BA. (**A**) Experimental setup. Juxtacellular labeling of principal neurons in the LA and BA of anesthetized mice (LAa: n=2; LAp: n=4; BAa: n=3, BAp: n=12; 21 PNs from 19 mice). (**B**) Dendritic reconstruction and Sholl analysis of *in vivo* filled example neurons from each subnucleus. Scale: 50 µm difference between the radii of concentric circles. (**C**) Scatter plot showing the total dendritic length of the *in vivo* filled principal neurons. (**D**) The dendritic length as a function of distance from the soma. (**E**) Number of dendritic branching points of 21 *in vivo* reconstructed neurons from each subnucleus. (**F**) A juxtacellularly filled example neuron with a reconstructed axonal and dendritic arborization. The dendrites are shown in black, and the axons in red innervating brain areas across different sections from the fundus of striatum (Br. -0.5 mm) to the subiculum (Br. -3.4 mm). The lateral subnucleus of the CEA (CEAl) is shown in green, while the medial and capsular subnuclei (CEAm+c) are shown in beige. Scale bar: 500 µm. The distance from Bregma (Br) is shown in mm. See the List of Abbreviations for the identification of brain areas.

Next, we compared the distribution of axonal collaterals of the reconstructed PNs in different amygdalar nuclei as well as the surrounding regions. An example PN shown in **Fig. 7F** demonstrates that its axons could be traced from the fundus of striatum (Bregma -0.5 mm) to the subiculum (Bregma -3.4 mm). Our analysis revealed characteristic axonal distributions for PNs located in the four subnuclei studied (**Fig. 8A**). The 2 LAa PNs had axons locally in the LA and BA and projected to the CEA and ventral part of the caudoputamen (CP, often referred to as the amygdalo-striatal area, Astria) (**Fig. 8B, C**). LAp PNs had more complex axonal arbors (**Fig. 8D**, **S7A**). They densely arborized locally within the LA, innervated the CP and projected either to the BAa or to the BAp. Anterior to the cell body, their axons reached the posterior insular cortex (AIp), whereas the posterior branches arborized in the amygdalo-piriform transition area (TR) (**Fig. 8D**). The axons of BAa PNs primarily ramified locally and in the BAp (**Fig. 8E**). They were observed in the medial, but not in the lateral nucleus of the CEA, an observation which is in line with our retrograde tracing results (**Fig. 6**, **S7A, B**). In contrast, the axon collaterals of BAp PNs were typically present in the lateral nucleus of the CEA in addition to the neighboring areas, including the BAa and TR (**Fig. 7F, 8F, G**). Interestingly, we observed axon collaterals of both BAa and BAp PNs in the LA, too (**Fig. 8**, **S7A, B**). Furthermore, 2 PNs were labeled in the BAp, the axons of which ramified extensively locally, and reached the different parts of the hippocampal formation (**Fig. S8**). These observations collectively show that, in addition to the subnucleus-restricted dendritic arbors of PNs in the LA and BA, their axons also arborize in a stereotypical manner, which may ensure a well-organized information flow within the BLA and the surrounding areas.

**Fig. 8.**
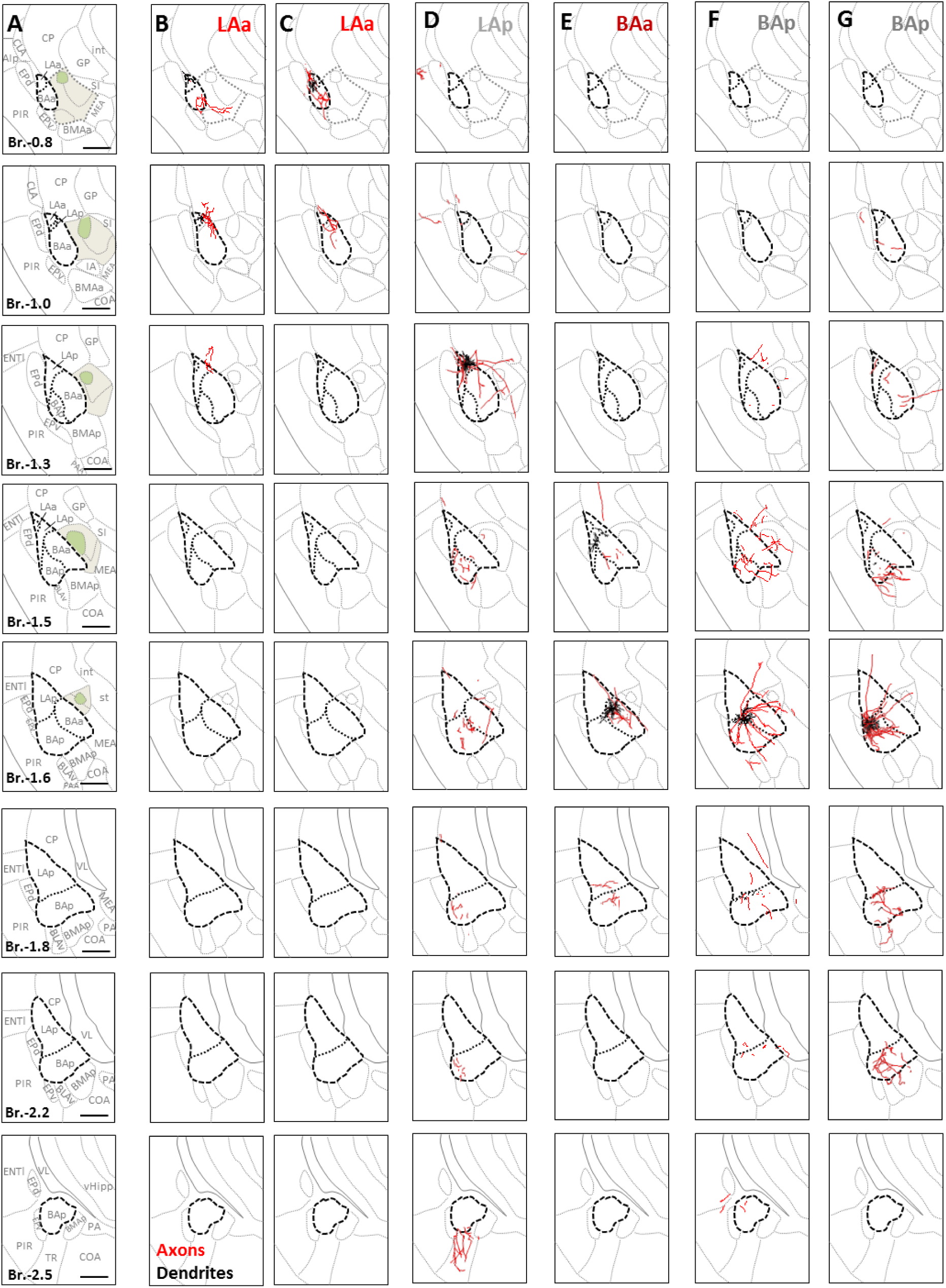
*In vivo* filled principal neurons in different LA and BA subnuclei with typical projection target sites. (**A**) Maps showing the LA and BA and the surrounding brain areas along 8 anteroposterior sections based on Allen Brain Mouse Brain Atlas (2011). The lateral subnucleus of the CEA (CEAl) is shown in green, while the medial and capsular subnuclei (CEAm+c) are shown in beige. (**B-C)** *In vivo* reconstructed small principal neurons in the LAa innervating the central amygdala (CEA) and the caudoputamen (CP). (**D**) A reconstructed large principal neuron located in the LAp. (**E**) A CeA-avoiding principal neuron with soma location in the BAa. (**F-G**) CEA-projecting principal neurons with soma location in the BAp. In each column, a single *in vivo* filled and reconstructed principal neuron is shown along the anteroposterior axis of the amygdala. Axons are shown in red and dendrites in black. Scale bar: 500 µm. Distance from Bregma (Br) is shown in mm. See the List of Abbreviations for the identification of brain areas.

## DISCUSSION

In this study we revealed that the DsRed expression driven by CCK promoter in BAC-CCK-DsRed mice (Mate et al., 2013; Rovira-Esteban et al., 2017) divides the LA and BA into two subnuclei. The borders of these CCK expression-defined areas do not completely align with the recently defined borders of LA and BA subnuclei: CCK+ BAa defined in our study corresponds mostly to the BLA.al and BLA.am, while the CCK-BAp matches the BLA.ac and BLAp subdomains defined by Hintiryan et al., 2021 (Hintiryan et al., 2021). Furthermore, the CCK+ BAa significantly but not completely overlaps with the BLA, but the CCK-BAp corresponds well to the BLP in the Mouse Brain Atlas (Paxinos & Franklin, 2004). The CCK+ LAa does not resemble any of the previous subdivisions in the mouse LA.

To assess whether these CCK expression-defined subunits can be considered as functional subnuclei, a comprehensive approach was employed: (1) the location of dendritic trees of LA and BA PNs was determined, (2) examples for selective innervation of LA and BA subnuclei were provided and (3) finally, distribution of PN populations with different projection areas were mapped relative to the CCK expression. We observed that the vast majority of dendritic trees (71-99%) were restricted to the subnucleus where the soma was located, which suggests independent information-receiving capabilities for PNs in the different subnuclei. In line with this conclusion, we found that prefrontal and insular cortices innervate LA and BA in a complementary manner; furthermore, in the medial portion along the anteroposterior axis of the LA and BA, the projections from the mPFC selectively terminate in the BAa, while the IC targets the LAp and BAp. Thus, CCK+ and CCK-BA subnuclei independently receive information from the two pathways examined. Based on this logic, CCK+ LAa PNs may receive different projections than CCK-LA PNs, a hypothesis that needs to be tested in future studies.

When amygdalar PN populations projecting to different areas were labeled by retrograde tracing, we found that PL-and DMS-projecting neurons were both exclusively located in the BAa, corroborating the earlier findings (Hintiryan et al., 2021; Hintiryan et al., 2016; Manoocheri & Carter, 2022; McGarry & Carter, 2017; O’Leary et al., 2020; Reppucci & Petrovich, 2016), including the simultaneous innervation of both brain areas (Fisher et al., 2020). Our findings are in accord with the previous results, showing that BA and LA PNs send collaterals to different downstream targets; for example, to the agranular insular cortex and nucleus accumbens (ACB) (Shinonaga et al., 1994) or to vHipp, mPFC and ACB (Beyeler et al., 2016). These data collectively support the logic of connectivity for cortical structures, indicating that if a given area projects to two distinct regions, then there will be neurons that project only to one of the target areas and there will be a set of neurons that project to both regions simultaneously (Lai et al., 2022; Shih & Chang, 2023). In contrast to the PNs projecting to the PL and DMS, the location of the PN population innervating the CEA is limited to the BAp, which is also consistent with recent findings (Beyeler et al., 2018; Massi et al., 2023). In the case of PN population projecting to the vHipp/SUB, no subnucleus-specificity was found, in accord with previous data (O’Leary et al., 2020). Although the rostral BAa→vHipp/SUB projection seems to be minimal based on our experiments, a recent study has provided support for the presence of CCK+ BA afferents in the vHipp/SUB, originating from the anterior part of the BA (Shen et al., 2019). We also observed a substantial number of LA PNs that were retrogradely labeled from the vHipp/SUB, which is in line with a previous tracing study (Petrovich et al., 2001). However, we cannot rule out the possibility that the spread of the retrograde tracers under our circumstances has reached the layer VI in the lateral entorhinal cortex, where LA PNs heavily project (Hintiryan et al., 2021), causing visualization of both LA PN populations projecting to the vHipp/SUB and/or lateral entorhinal cortex.

Our results show that while PL-and DMS-projecting PNs are predominantly in the CCK+ area, vHipp/SUB-projecting PNs may be composed of CCK+ and CCK-subpopulations, and CEA-projecting neurons are mostly in the CCK-subregions. The BA→PL pathway has a prominent role in the regulation of reward-seeking and fear-related behaviors (Burgos-Robles et al., 2017; Felix-Ortiz et al., 2016; Sotres-Bayon et al., 2012), as well as in social behaviors (Kietzman et al., 2022; Scheggia et al., 2022), whereas the BA→DMS connection may play a role in controlling obsessive-compulsive behavior (Yoon et al., 2023). The BLA→vHipp/SUB projection plays a role in coordinating social (Felix-Ortiz & Tye, 2014) and anxiety-related behaviors (Felix-Ortiz et al., 2013). As DsRed signal varies in vHipp/SUB-projecting BLA PNs, it may be worthwhile to separately examine the roles of CCK+ and CCK-PN populations in amygdala-controlled processes in future studies. The BAp→CEA innervation also contributes to the control of anxiety-like behavior (Tye et al., 2011) as well as to learning and expression of conditioned fear (Massi et al., 2023). These previous findings collectively underscore the hub-like nature of the BLA that channels the information flow to distinct downstream regions depending on the behavioral challenges and internal states.

*In vitro* and *in vivo* cell reconstructions were conducted to morphologically characterize and compare the properties of PNs of the different LA and BA subnuclei and reveal their local axonal arbors. Based on the properties of dendritic arborizations, the LAa PNs were significantly smaller and had less ramified dendrites than the LA PNs, which was also confirmed by their electrophysiological properties and supported by *in vivo* single-cell labeling. The presence of such small PNs in the LA has not been reported earlier (Duvarci & Pare, 2007; Faber et al., 2001; Rainnie et al., 1993). Although previous studies obtained in rats and monkeys suggest that BA subdivisions can be distinguished based on cell size into parvo-, intermediate-, and magnocellular parts (Pitkanen & Amaral, 1998; Savander et al., 1995), we found no obvious indication for cell size differences in the mouse BA.

Based on previous results (Beyeler et al., 2018; Hintiryan et al., 2021; Pitkanen, 2000; Sah et al., 2003), both LA and BA subnuclei receive distinct sets of inputs that, after processing, transmit to separate as well as overlapping output areas. LA combines cortical inputs primarily from higher order auditory cortices, temporal association cortex, and the IC with inputs from the posterior thalamic nuclei and sends collaterals to the subnuclei of the BA and TR in addition to the ventral part of the CP (i.e., to the Astria) and CEA. In contrast, the main input-output cortical structure of the BAa is the mPFC. Importantly, BAa PNs send axons remotely to the mPFC and DMS and locally to BAp circuits and the medial part of the CEA, but not to its lateral part. BAp neurons receiving afferents from the IC heavily project to both parts of the CEA in addition to the BAa and TR. Thus, both the lateral nucleus of the CEA and TR receive inputs from LA and BAp, but not from the BAa, whereas the output site of the CEA, the medial nucleus is innervated by LA, BAa and BAp. In addition to cortical inputs, BA receives excitatory innervation from the midline thalamic nuclei (MT) (Amir et al., 2019; Matyas et al., 2018; Turner & Herkenham, 1991), which also parcel the BA. BAa receives inputs primarily from the central medial nucleus of thalami, which does not target the CEA, whereas the paraventricular nucleus of thalami innervates both the BAp and CEA (Amir et al., 2019; Turner & Herkenham, 1991). Interestingly, we observed many axon collaterals of *in vivo* labeled BA PNs in the LA (**Fig. 8E-G**), an observation which suggests that bidirectional communication between the LA and BA may exist that has not been recognized earlier. Future studies should clarify the role of the BA to LA connectivity.

In summary, the functional subnuclei in the LA and BA – defined by CCK expression and restricted dendritic arbors of PNs – receive a specific combination of afferents and project to well-defined intra-and extra-amygdalar areas. The complex organization of inputs to the subnuclei and their outputs may ensure the critical combination and coding of information, necessary for predictive processing that is a key cortical function supporting adaptive behavior (Keller & Mrsic-Flogel, 2018; Lee et al., 2021).

## MATERIALS AND METHODS

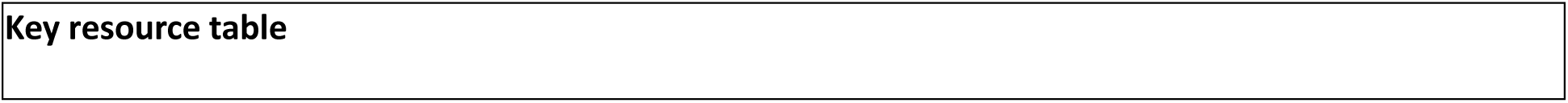
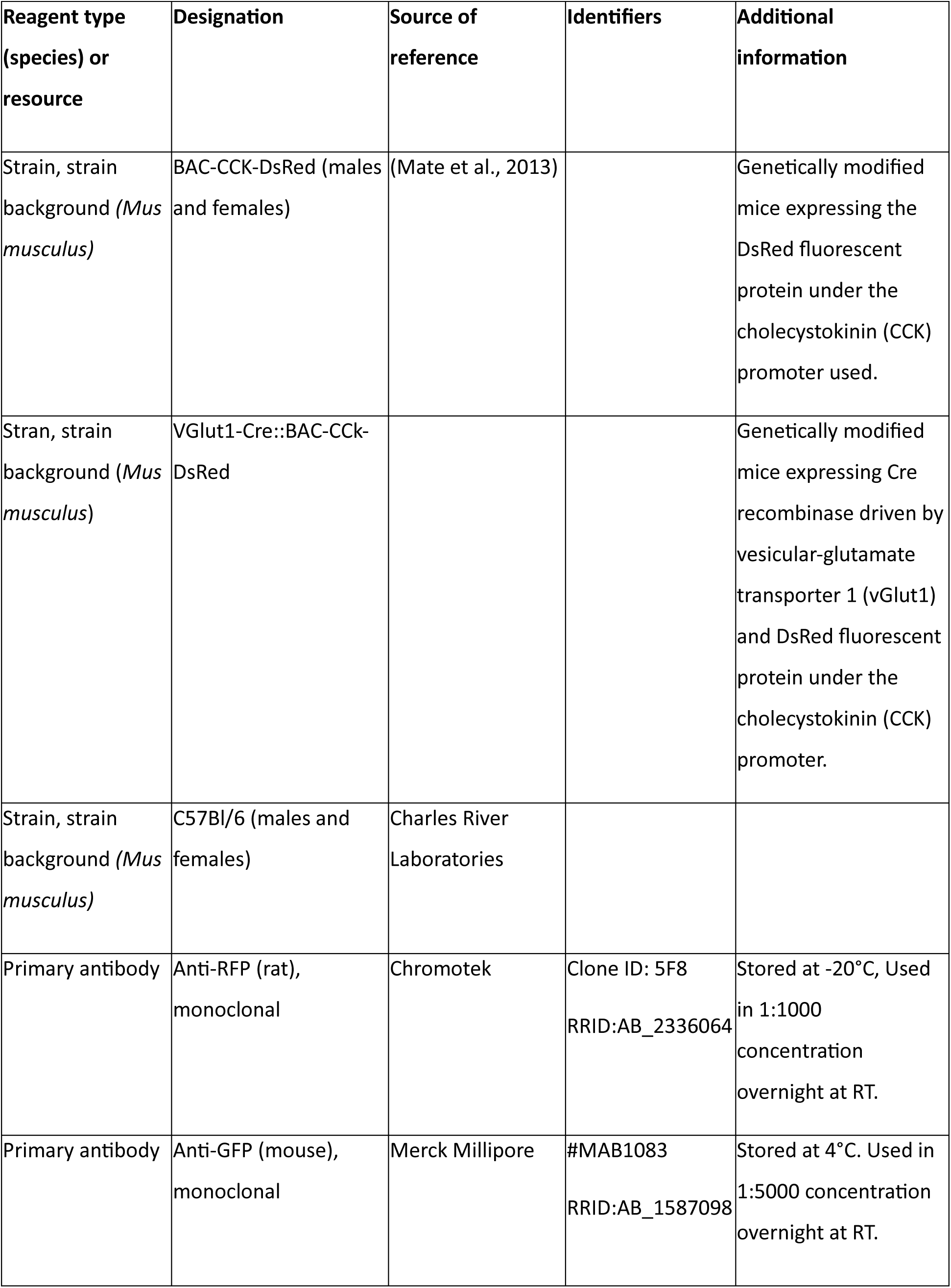
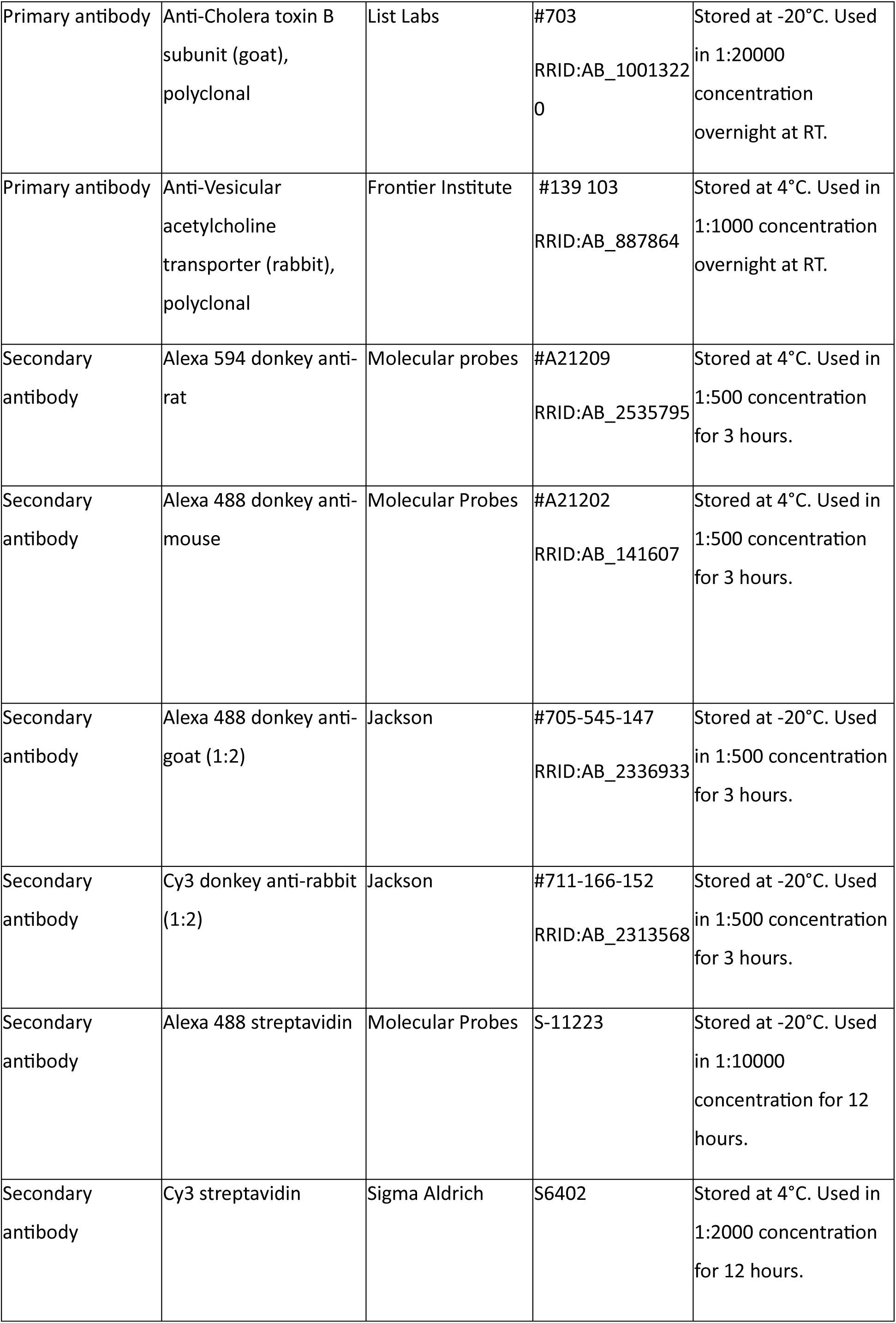
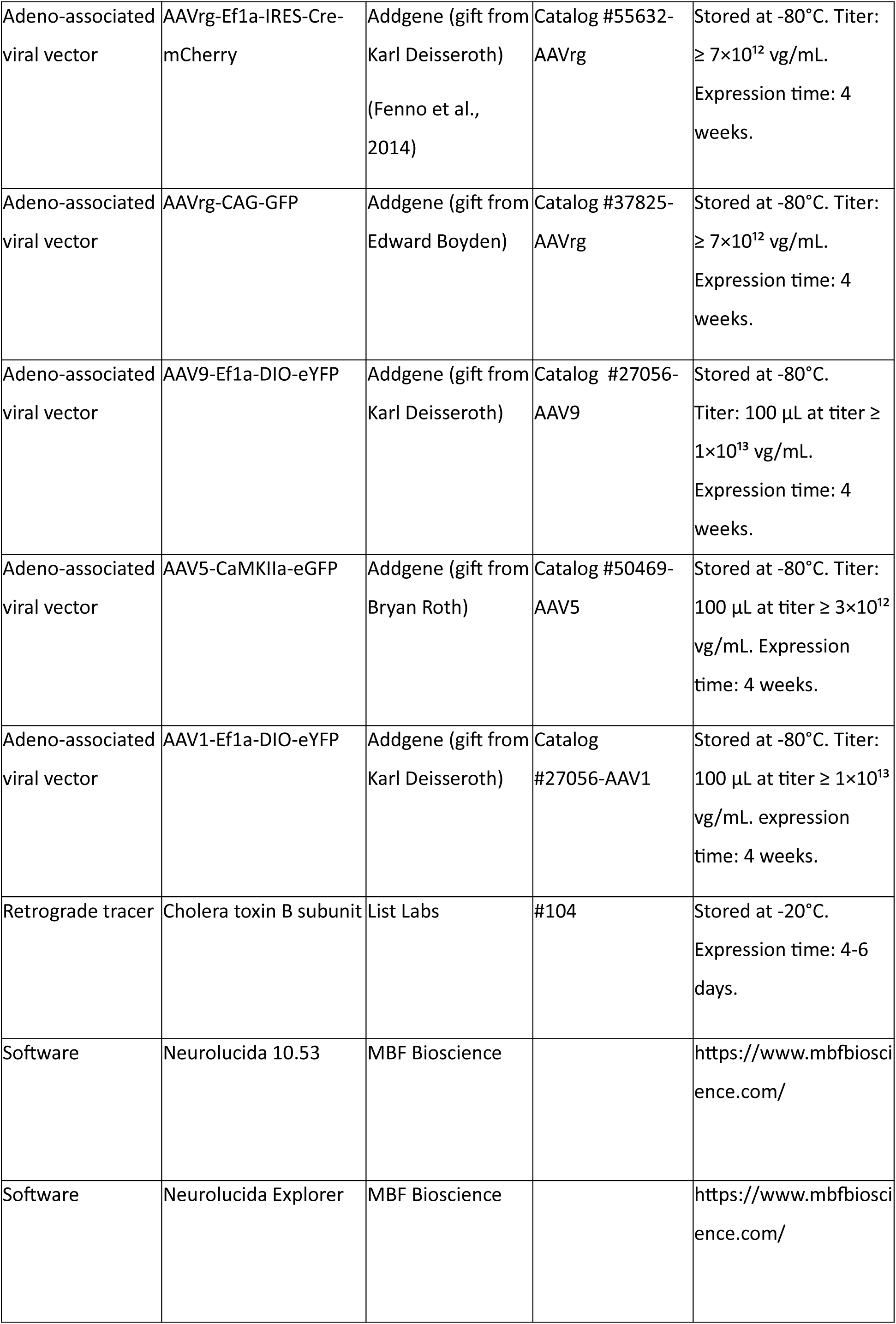
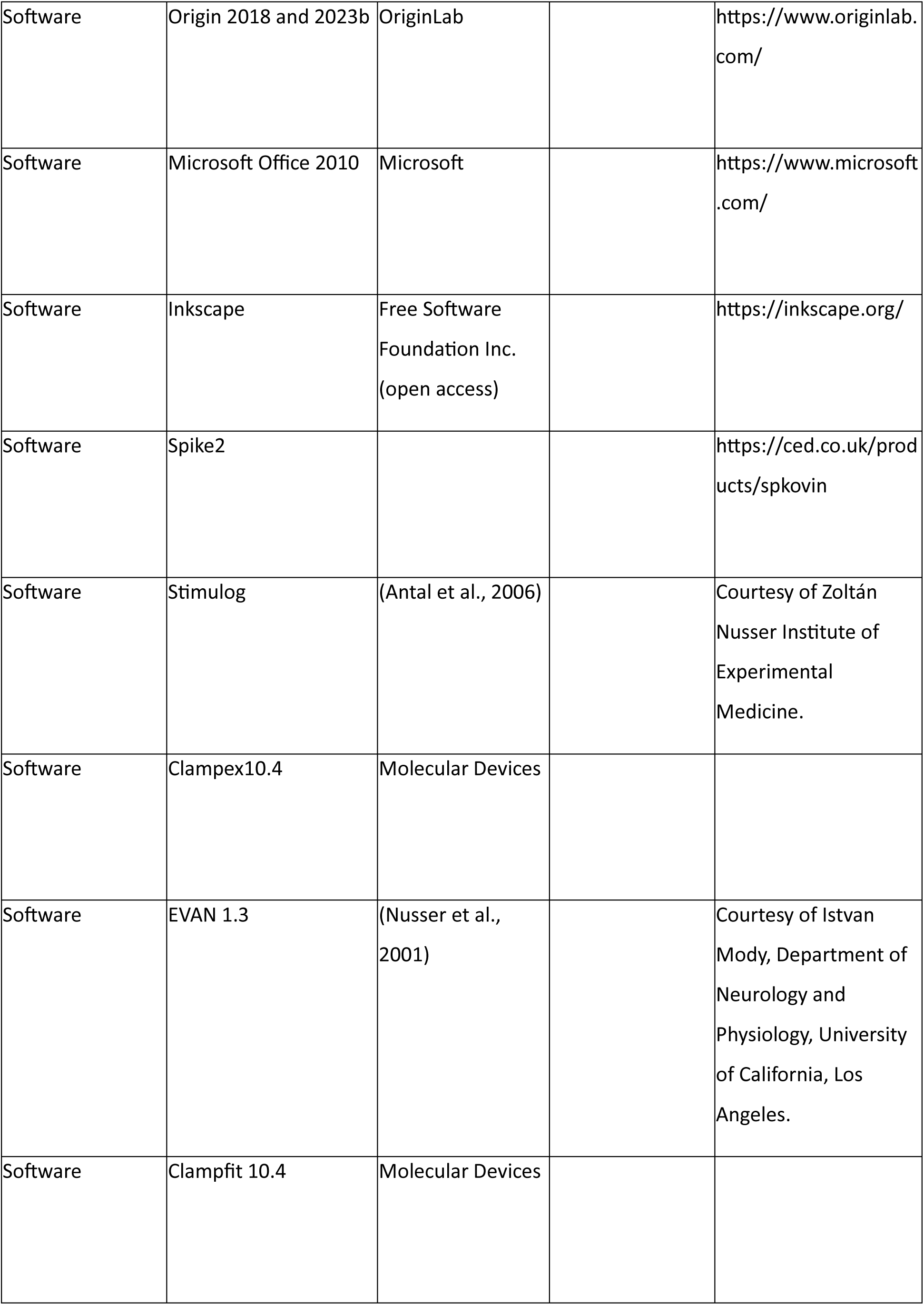

|  |
| --- |
| Key resource table |

| Reagent type<br>(species) or<br>resource | Designation | Source of<br>reference | Identifiers | Additional<br>information |
| --- | --- | --- | --- | --- |
| Strain, strain<br>background (*Mus*<br>*musculus*) | BAC-CCK-DsRed (males<br>and females) | (Mate et al., 2013) |  | Genetically modified<br>mice expressing the<br>DsRed fluorescent<br>protein under the<br>cholecystokinin (CCK)<br>promoter used. |
| Stran, strain<br>background (*Mus*<br>*musculus*) | VGlut1-Cre::BAC-Cck-DsRed |  |  | Genetically modified<br>mice expressing Cre<br>recombinase driven by<br>vesicular-glutamate<br>transporter 1 (vGlut1)<br>and DsRed fluorescent<br>protein under the<br>cholecystokinin (CCK)<br>promoter. |
| Strain, strain<br>background (*Mus*<br>*musculus*) | C57Bl/6 (males and<br>females) | Charles River<br>Laboratories |  |  |
| Primary antibody | Anti-RFP (rat),<br>monoclonal | Chromotek | Clone ID: 5F8<br>RRID:AB_2336064 | Stored at -20°C, Used<br>in 1:1000<br>concentration<br>overnight at RT. |
| Primary antibody | Anti-GFP (mouse),<br>monoclonal | Merck Millipore | #MAB1083<br>RRID:AB_1587098 | Stored at 4°C. Used in<br>1:5000 concentration<br>overnight at RT. |
| Primary antibody | Anti-Cholera toxin B subunit (goat), polyclonal | List Labs | #703<br>RRID:AB_10013220 | Stored at -20°C. Used in 1:20000 concentration overnight at RT. |
| Primary antibody | Anti-Vesicular acetylcholine transporter (rabbit), polyclonal | Frontier Institute | #139 103<br>RRID:AB_887864 | Stored at 4°C. Used in 1:1000 concentration overnight at RT. |
| Secondary antibody | Alexa 594 donkey anti-rat | Molecular probes | #A21209<br>RRID:AB_2535795 | Stored at 4°C. Used in 1:500 concentration for 3 hours. |
| Secondary antibody | Alexa 488 donkey anti-mouse | Molecular Probes | #A21202<br>RRID:AB_141607 | Stored at 4°C. Used in 1:500 concentration for 3 hours. |
| Secondary antibody | Alexa 488 donkey anti-goat (1:2) | Jackson | #705-545-147<br>RRID:AB_2336933 | Stored at -20°C. Used in 1:500 concentration for 3 hours. |
| Secondary antibody | Cy3 donkey anti-rabbit (1:2) | Jackson | #711-166-152<br>RRID:AB_2313568 | Stored at -20°C. Used in 1:500 concentration for 3 hours. |
| Secondary antibody | Alexa 488 streptavidin | Molecular Probes | S-11223 | Stored at -20°C. Used in 1:10000 concentration for 12 hours. |
| Secondary antibody | Cy3 streptavidin | Sigma Aldrich | S6402 | Stored at 4°C. Used in 1:2000 concentration for 12 hours. |
| Adeno-associated viral vector | AAVrg-Ef1a-IRES-Cre-mCherry | Addgene (gift from Karl Deisseroth) (Fenno et al., 2014) | Catalog #55632-AAVrg | Stored at -80°C. Titer: $\geq 7 \times 10^{12}$ vg/mL. Expression time: 4 weeks. |
| Adeno-associated viral vector | AAVrg-CAG-GFP | Addgene (gift from Edward Boyden) | Catalog #37825-AAVrg | Stored at -80°C. Titer: $\geq 7 \times 10^{12}$ vg/mL. Expression time: 4 weeks. |
| Adeno-associated viral vector | AAV9-Ef1a-DIO-eYFP | Addgene (gift from Karl Deisseroth) | Catalog #27056-AAV9 | Stored at -80°C. Titer: 100 μL at titer $\geq 1 \times 10^{13}$ vg/mL. Expression time: 4 weeks. |
| Adeno-associated viral vector | AAV5-CaMKIIa-eGFP | Addgene (gift from Bryan Roth) | Catalog #50469-AAV5 | Stored at -80°C. Titer: 100 μL at titer $\geq 3 \times 10^{12}$ vg/mL. Expression time: 4 weeks. |
| Adeno-associated viral vector | AAV1-Ef1a-DIO-eYFP | Addgene (gift from Karl Deisseroth) | Catalog #27056-AAV1 | Stored at -80°C. Titer: 100 μL at titer $\geq 1 \times 10^{13}$ vg/mL. expression time: 4 weeks. |
| Retrograde tracer | Cholera toxin B subunit | List Labs | #104 | Stored at -20°C. Expression time: 4-6 days. |
| Software | Neurolucida 10.53 | MBF Bioscience |  | https://www.mbfbioscience.com/ |
| Software | Neurolucida Explorer | MBF Bioscience |  | https://www.mbfbioscience.com/ |
| Software | Origin 2018 and 2023b | OriginLab |  | https://www.originlab.com/ |
| Software | Microsoft Office 2010 | Microsoft |  | https://www.microsoft.com/ |
| Software | Inkscape | Free Software Foundation Inc. (open access) |  | https://inkscape.org/ |
| Software | Spike2 |  |  | https://ced.co.uk/products/spkovin |
| Software | Stimulog | (Antal et al., 2006) |  | Courtesy of Zoltán Nusser Institute of Experimental Medicine. |
| Software | Clampex10.4 | Molecular Devices |  |  |
| Software | EVAN 1.3 | (Nusser et al., 2001) |  | Courtesy of Istvan Mody, Department of Neurology and Physiology, University of California, Los Angeles. |
| Software | Clampfit 10.4 | Molecular Devices |  |  |

### Subjects

Adult mice (median: 124.5, interquartile range: 138) of both sexes were used for the experiments. BAC-CCK-DsRed (Mate et al., 2013) transgenic line expressing DsRed fluorescent protein under cholecystokinin (CCK) promoter was used for *in vitro* measurements (n=37) and to label midline thalamic afferents reaching the amygdala (n=2). VGlut1-Cre::BAC-CCk-DsRed animals expressing Cre driven by vesicular-glutamate transporter type 1 (vGlut1) promoter and DsRed fluorescent protein under CCK promoter were used for anterograde tracing experiments labeling cortical afferents (n=2-3). C57Bl/6 (Charles River) wild type mice were used for retrograde tracing (n=12), collateral labeling (n=2-2) and *in vivo* juxtacellular (n=55) experiments. Animals were kept in groups of 2-4 mice in transparent Plexiglas cages, under 12-hour light/dark cycle. The constant temperature and humidity of the environment were controlled. Mice had *ad libitum* access to food and water.

Experiments were approved by the Committee of the Scientific Ethics of Animal Research (22.1/360/3/2011) and all procedures involving animals were performed according to methods approved by Hungarian legislation (1998 XXVIII. section 243/1998, renewed in 40/2013.) and institutional guidelines of ethical code. All procedures complied with the European convention for the protection of vertebrate animals used for experimental and other scientific purposes (Directive 86/609/CEE and modified according to the Directives 2010/63/EU). Every effort was taken to minimize animal suffering and the number of animals used.

### Antero-and retrograde tracing

Mice were anesthetized with isoflurane or 125 mg/kg ketamine and 5 mg/kg xylazine (all from Medicus Partner) and secured in a stereotaxic frame. Anteroposterior (AP) and mediolateral (ML) coordinates were measured from the Bregma, while dorsoventral (DV) coordinate from the surface of dura mater. Small holes over the injection sites were created using a dental drill (Foredom).

#### Anterograde tracing experiments

VGlut1-Cre::BAC-CCK-DsRed animals were injected with AAV9-Ef1a-DIO-eYFP (Addgene 27056-AAV; titer: 100 µL at titer ≥ 1×10¹³ vg/mL) virus bilaterally, targeting the: medial prefrontal cortex (mPFC; AP: 1.5 and 2.0 mm; ML: 0.5 mm and DV: 2.0 and 1.5 mm) and insular cortex (IC; AP: 2.0 mm; ML: 2.5 mm; DV: 2.0 mm). AAV5-CAMKII-eGFP (Addgene 50469-AAV5; 100 µL at titer ≥ 3×10¹² vg/mL) was used to inject the midline thalamus (MT; AP: -1.8 mm; ML: 0.4 mm and DV: 3.2 mm) using Nanoject III Programmable Nanoliter Injector (Drummond Scientific Company).

#### Retrograde tracing experiments

AAVrg-Ef1a-IRES-Cre-mCherry (Addgene, 55632-AAVrg; titer: ≥ 7×10¹² vg/mL; 100 nl/injection site), AAVrg-CAG-GFP (Addgene, 37825-AAVrg; titer: ≥ 7×10¹² vg/mL; 100 nl/injection site) or 0.5% Cholera toxin B subunit (CTB; List Labs; 25 nl/injection site) were used aiming one of the following target sites uni-or bilaterally: prelimbic cortex (PL; AP: 1.78 and 2.1 mm; ML: 0.3 mm; DV: 1.0 mm; n=2), dorsomedial striatum (DMS; AP: 0.62 and 1.1 mm; ML: 1.3 mm; DV: 2.3 mm; n=2), and subiculum (S; AP: -3.5 and -3.8 mm; ML: 3.4 mm; DV: 4.1 mm). In the case of experiments that aimed to co-label two neuronal populations and the overlap between them (n=2), the viral vectors were injected to the PL and DMS, respectively. Tracers were applied to the brain through a glass pipette (ID = 0.530 mm ± 25 um, OD 1.14 mm, World Precision Instruments). The flow rate of the injections was 1 nl/sec. Due to its small size, the central amygdala (CEA; AP: -1.6 mm; ML: 2.6 mm; DV: 4.1 mm) was injected iontophoretically (2 or 5 µA pulses with 2/2 s on/off duty cycle for 7 or 10 minutes, respectively) with CTB using a Drummond Recording Nanoject II (Drummond Scientific).

#### Visualizing axon collaterals

To label the axon collaterals of the DMS-and PL-projecting amygdalar PNs an intersectional approach was used: AAVrg-Ef1a-IRES-Cre-mCherry was injected to the PL or DMS at two anteroposterior coordinates, together with AAV1-Ef1a-DIO-eYFP (Addgene, 27056-AAV1; titer: 100 µL at titer ≥ 1×10¹³ vg/mL) administered to the BA (AP: -1.5 mm; ML: 3.1 mm and DV: 4.25 mm).

Processing brain tissue. After 4-5 days (in case of CTB injections) or 4 weeks (in case of AAV injections) of survival, mice were transcardially perfused with 4% paraformaldehyde (PFA; Sigma-Aldrich) in 0.1 M phosphate-buffer (PB), pH 7.4, for 40-50 minutes, with a flow rate of 3 ml/min. On the same day, the brains were cut into 100 µm thick slices using a Vibratome (Leica VT1000S). Finally, the slices were stored in 0.1M PB containing 0.05% Na-azide at a temperature of 4°C until further processing.

### *In vitro* recording and labeling

For preparing acute brain slices, mice were deeply anesthetized with isoflurane and decapitated. The brain was quickly removed and placed into ice-cold solution, containing (in mM): 252 sucrose, 2.5 KCl, 26 NaHCO_3_, 0.5 CaCl_2_, 5 MgCl_2_, 1.25 NaH_2_PO4, 10 glucose, bubbled with 95% O_2_/5% CO_2_ (carbogen gas). Coronal or horizontal slices of 200-350 μm thickness containing the LA or BA were prepared with a Leica VT1000S or VT1200S vibratome and kept in an interface-type holding chamber containing ACSF at 36°C that gradually cooled down to room temperature. ACSF contained the followings (in mM): 126 NaCl, 2.5 KCl, 1.25 NaH_2_PO_4_, 2 MgCl_2_, 2 CaCl_2_, 26 NaHCO_3_, and 10 glucose, bubbled with carbogen gas.

After at least 1 hour incubation, slices were transferred to a submerged type recording chamber perfused with 32°C ACSF with approximately 2–2.5 ml/min flow rate. Recordings were performed under visual guidance using differential interference contrast microscopy (Olympus BX61W or Nikon FN-1) using 40x water dipping objective. Neurons expressing DsRed were visualized with the aid of a mercury arc lamp or a monochromator (Till Photonics) and detected with a CCD camera (Hamamatsu Photonics or Andor Zyla). Patch pipettes (4–7 MΩ) for whole-cell recordings were pulled from borosilicate capillaries with inner filament (thin walled, OD 1.5) using a DMZ-Universal Puller (Zeitz Instruments) or using a P1000 pipette puller (Sutter Instruments). In whole-cell recordings the patch pipette contained a K-gluconate-based intrapipette solution containing the following (in mM): 110 K-gluconate, 4 NaCl, 2 Mg-ATP, 20 HEPES, 0.1 EGTA, 0.3 GTP (sodium salt), and 10 phosphocreatine adjusted to pH 7.3 using KOH, with an osmolarity of 290 mOsm/L and additional 0.2% biocytin.

Recordings were performed with a Multiclamp 700B amplifier (Molecular Devices), low-pass filtered at 3 kHz, digitized at 10 kHz, recorded with an in-house data acquisition and stimulus software (Stimulog, courtesy of Zoltán Nusser, Institute of Experimental Medicine) or Clampex 10.4 (Molecular Devices), and were analyzed with EVAN 1.3 (courtesy of Istvan Mody, Department of Neurology and Physiology, University of California, Los Angeles), Clampfit 10.4 (Molecular Devices), and OriginPro 2018 (OriginLab). Recordings were not corrected for junction potential. To test the firing characteristics, neurons were injected with 800-ms–long hyperpolarizing and depolarizing square current pulses with increasing amplitudes from -100 to 600 pA. Principal neurons were identified based on their broad action potential waveform, accommodating firing pattern, and slow after-hyperpolarization in addition to their *post-hoc* identified morphological appearance.

### *In vivo* recording and labeling

The anesthesia was induced with isoflurane and maintained with 125 mg/kg ketamine and 5 mg/kg xylazine. A cranial window was opened above the amygdala and a glass electrode containing 3% neurobiotin (Vector Laboratories) in 0.5M NaCl was lowered to the target depth (4.8 mm). After the amygdala has been reached, the pipette was moved forward slowly (1 μm/sec) until contacting a neuron. Using positive current steps, the neuron was loaded with neurobiotin tracer as described in Pinault (1996) (Pinault, 1996). After successful labeling, the glass electrode was withdrawn from the brain. Then, animals were perfused as described above. Brains were cut to 80-150 µm thick slices and stored in 0.1 M PB containing 0.05% Na-azide at a temperature of 4°C until further processing.

### Immunohistochemistry

#### Tracing experiments

Sections underwent an initial washing process in 0.1 M PB consisting of 3 washing steps lasting 10 minutes each. Then, the sections were blocked for 30 minutes using a solution of 10% Normal Donkey Serum (NDS, Vector Laboratories) and 0.5% Triton X-100 (Acros Organics) diluted in 0.1 M PB. This was followed by treatment with a solution containing 2% NDS, 0.5% Triton-X, 0.05% Na-azide and primary antibodies diluted in 0.1 M PB for 1 night at RT. In virus tracing experiments, mouse anti-GFP (mGFP; Molecular Probes; 1:5000) and rat anti-RFP (ratRFP; Chromotek; 1:1000) were used to enhance the signal of GFP and RFP, respectively. In CTB-tracing experiments, goat anti-CTB (gCTB; List Labs; 1:20000) was applied to detect the retrogradely labelled cells. Then, the sections were thoroughly washed (3-4 washing steps, 15 min/wash) and incubated for 3 hours in a solution comprising of 1% NDS and the secondary antibodies diluted in 0.1 M PB. The secondary antibodies were the followings: Alexa 488 donkey anti-mouse (A488 DAM; Molecular Probes; 1:500) and Alexa 594 donkey anti-rat (A594 DARat; Molecular Probes; 1:500) to enhance the visualization of mGFP and ratRFP, respectively; and Alexa 488 donkey anti-goat (A488 DAG; Jackson; 1:500) to reveal gCTB. Vesicular acetylcholine transporters (VAChT) were visualized using rabbit anti-VAChT primary antibody (Frontier Institute, 1:1000), aiding the identification of borders between BA and LA (Arvidsson et al., 1997; Vereczki et al., 2021). Finally, following several washes in PB, sections were mounted on glass slides in Vectashield (Vector Laboratories) or ProLong Diamond Antifade Mountant (Diagnosticum).

#### Visualization of in vitro and in vivo filled neurons

The visualization process of filled neurons, both *in vitro* and *in vivo*, involved the following steps: after the initial washing in tris-buffered saline (TBS), the slices were incubated for 12 hours in a solution containing 0.5% Triton-X and fluorophore-conjugated streptavidin (Alexa488-SA, Molecular Probes; 1:10000 or Cy3-SA; Sigma-Aldrich; 1:2000). For *in vivo* filled neurons VAChT was visualized to help the identification of borders between the BA and LA. This step was followed by multiple washing in TBS and incubation in a solution containing Cy3 donkey anti rabbit (Cy3 DAR; Jackson; 1:500) secondary antibody diluted in TBS. Finally, the slices underwent thorough washing, 3 times in TBS and 3 more times in 0.1 M PB (10 min/wash) and then were mounted on glass slides in Vechtashield.

#### Confocal microscopy

Multichannel fluorescent images at high resolution were acquired with Nikon C2 confocal laser scanning microscope (Nikon Europe, Amsterdam, the Netherlands) in channel series mode. Plan Apo VC 20x (NA=0.75) objective was used to take images for 3D reconstructions (z step size: 1 or 2 µm; xy: 0.61-0.63 µm/px) and to determine the location of retrogradely labelled neurons in the LA and BA (1 focal plane/section; xy: 0.63 µm/px). Coronal and horizontal amygdala-sections of BAC-CCK-DsRed animals, mPFC and IC inputs reaching the LA and BA, and the collaterals of the PL-and DMS-projecting neurons were captured with Plan Fluor 10x (NA=0.3) objective (1 focal plane/section; xy: 0.93-1.23 µm/px).

#### Reconstruction and analysis

Principal neuronal identity was confirmed *post hoc* by their characteristic spiny dendrites and morphological appearance. Reconstruction and analysis of *in vitro* and *in vivo* filled neurons was performed using Neurolucida 10.53 software (MBF Bioscience) (1 PN was reconstructed from 9-37 sections). The properties of axonal and dendritic arbors, including lengths, node numbers, and Sholl analysis, were determined using Neurolucida Explorer (MBF Bioscience). Tissue shrinkage along the x and y axes was quantified by comparing the distance between brain regions in the same sections before and after the mounting process. Shrinkage along the z-axis was calculated based on the difference in section thickness before and after mounting. Finally, correction values were the followings: xy: 1 and z: 2.5 for *in vitro* slices and xyz: 1.185 for *in vivo* sections (using Plan Apo VC 20x objective). The shrinkage caused by the fixation processes of the slices was not considered. The borders of the amygdala and the surrounding brain areas were drawn based on VAChT staining and the Allen Mouse Brain Atlas (Allen Institute 2011). In this study, the nomenclature of the Allen Brain Atlas was used.

The analysis of retrograde injections involved marking the location of retrogradely labeled neurons in a single focal plane in the LA and BA using Neurolucida. The neuron populations projecting to the PL, DMS, CEA or vHipp/SUB were shown in amygdalar sections from different planes along the AP axis.

Schematic diagrams illustrating brain areas, brain sections with injections sites, and experiments were created using Inkscape, an open-access program provided by the Free Software Foundation.

#### Statistics

To assess whether the data were drawn from a normal distribution was tested using the Shapiro-Wilk test. Since the datasets typically did not exhibit a normal distribution, comparisons between certain groups of anatomical and electrophysiological data were conducted using the Kruskal-Wallis non-parametric test. For *post hoc* analysis Dunn’s test was used. All statistics were performed using Origin 8.6 or 9.2 240 (Northampton, MA). Exact p values were indicated when p was higher than 0.001 considering the rounding rules. Data are presented as mean ± SEM, unless indicated otherwise. On box charts the mean (small open square), the median (continuous line within the box), the interquartile range (box) and the 5-95% values (ends of whiskers bar) are plotted. In the scatter plots, each symbol represents individual data points. Due to the small number of elements no statistical test was used for the representation of the *in vivo* filled principal neurons.

## ACKNOWLEDGEMENTS

We acknowledge financial support from the HUN-REN Hungarian Research Network, Hungarian Brain Research Program (2017-1.2.1-NKP-2017-00002) and National Research, Development and Innovation Office (K131893). The authors are grateful to Bence Barabás, Erzsébet Gregori, Éva Krizsán and Péter Laár for their excellent technical assistance. We also thank László Barna and Pál Vági, the Nikon Microscopy Center at the Institute of Experimental Medicine, Nikon Austria GmbH, and Auro-Science Consulting, Ltd., for kindly providing microscopy support.

## LIST OF ABBREVIATIONS

AAA: anterior amygdalar area
AIp: agranular insular area, posterior part
AP: anteroposterior
BAa: basal amygdalar nucleus, anterior part
BAp: basal amygdalar nucleus, posterior part
BLAv: basolateral amygdalar nucleus, ventral part
BMAa: basomedial amygdalar nucleus, anterior part
BMAp: basomedial amygdalar nucleus, posterior part
Br.: Bregma
CEA: central amygdalar nucleus
CEAl: central amygdalar nucleus, lateral part
CEAm+c: central amygdalar nucleus, medial and capsular part
CLA: claustrum
COA: cortical amygdalar area
CP: caudoputamen
cpd: cerebral peduncle
CTB: Cholera toxin B subunit
ctx: cortex
DMS: dorsomedial striatum
DV: dorsoventral
ec: external capsule
ENTl: entorhinal area, lateral part
ENTmv: entorhinal area, medial part, ventral zone
EP nucl.: endopiriform nucleus
EPd: endopiriform nucleus, dorsal part
EPv: endopiriform nucleus, ventral part
fi: fimbria
FS: fundus of striatum
GP: globus pallidus
IA: Intercalated amygdalar nucleus
int: internal capsule
LA: lateral amygdalar nucleus
LAa: lateral amygdalar nucleus, anterior part
LAp: lateral amygdalar nucleus, posterior part
MEA: medial amygdalar nucleus
ML: mediolateral
opt: optic tract
PA: posterior amygdalar nucleus
PAA: piriform-amygdalar area
PFC: prefrontal cortex
PIR: piriform cortex
PL: prelimbic cortex
RT: reticular nucleus of the thalamus
SI: substania innominata
st: stia terminalis
SUBv: subiculum, ventral part
TR: postpiriform transition area
vHipp: ventral hippocampus
vHipp/SUB: ventral hippocampus/subiculum
VL: lateral ventricle
ZI: zona incerta

## REFERENCES

Ahmed, N., Headley, D. B., & Pare, D. (2021). Optogenetic study of central medial and paraventricular thalamic projections to the basolateral amygdala. J Neurophysiol, 126(4), 1234–1247. 10.1152/jn.00253.2021

Amir, A., Pare, J. F., Smith, Y., & Pare, D. (2019). Midline thalamic inputs to the amygdala: Ultrastructure and synaptic targets. J Comp Neurol, 527(5), 942–956. 10.1002/cne.24557

Antal, M., Eyre, M., Finklea, B., & Nusser, Z. (2006). External tufted cells in the main olfactory bulb form two distinct subpopulations. Eur J Neurosci, 24(4), 1124–1136. 10.1111/j.1460-9568.2006.04988.x

Arvidsson, U., Riedl, M., Elde, R., & Meister, B. (1997). Vesicular acetylcholine transporter (VAChT) protein: a novel and unique marker for cholinergic neurons in the central and peripheral nervous systems. J Comp Neurol, 378(4), 454–467. https://www.ncbi.nlm.nih.gov/pubmed/9034903

Beyeler, A., Chang, C. J., Silvestre, M., Leveque, C., Namburi, P., Wildes, C. P., & Tye, K. M. (2018). Organization of Valence-Encoding and Projection-Defined Neurons in the Basolateral Amygdala. Cell Rep, 22(4), 905–918. 10.1016/j.celrep.2017.12.097

Beyeler, A., Namburi, P., Glober, G. F., Simonnet, C., Calhoon, G. G., Conyers, G. F., Luck, R., Wildes, C. P., & Tye, K. M. (2016). Divergent Routing of Positive and Negative Information from the Amygdala during Memory Retrieval. Neuron, 90(2), 348–361. 10.1016/j.neuron.2016.03.004

Bowers, M. E., & Ressler, K. J. (2015). Interaction between the Cholecystokinin and Endogenous Cannabinoid Systems in Cued Fear Expression and Extinction Retention. Neuropsychopharmacology, 40(3), 688–700. 10.1038/npp.2014.225

Burgos-Robles, A., Kimchi, E. Y., Izadmehr, E. M., Porzenheim, M. J., Ramos-Guasp, W. A., Nieh, E. H., Felix-Ortiz, A. C., Namburi, P., Leppla, C. A., Presbrey, K. N., Anandalingam, K. K., Pagan-Rivera, P. A., Anahtar, M., Beyeler, A., & Tye, K. M. (2017). Amygdala inputs to prefrontal cortex guide behavior amid conflicting cues of reward and punishment. Nat Neurosci, 20(6), 824–835. 10.1038/nn.4553

Duvarci, S., & Pare, D. (2007). Glucocorticoids enhance the excitability of principal basolateral amygdala neurons. J Neurosci, 27(16), 4482–4491. 10.1523/JNEUROSCI.0680-07.2007

Erlich, J. C., Bush, D. E., & Ledoux, J. E. (2012). The role of the lateral amygdala in the retrieval and maintenance of fear-memories formed by repeated probabilistic reinforcement. Front Behav Neurosci, 6, 16. 10.3389/fnbeh.2012.00016

Faber, E. S., Callister, R. J., & Sah, P. (2001). Morphological and electrophysiological properties of principal neurons in the rat lateral amygdala in vitro. J Neurophysiol, 85(2), 714–723. 10.1152/jn.2001.85.2.714

Felix-Ortiz, A. C., Beyeler, A., Seo, C., Leppla, C. A., Wildes, C. P., & Tye, K. M. (2013). BLA to vHPC Inputs Modulate Anxiety-Related Behaviors. Neuron, 79(4), 658–664. 10.1016/j.neuron.2013.06.016

Felix-Ortiz, A. C., Burgos-Robles, A., Bhagat, N. D., Leppla, C. A., & Tye, K. M. (2016). Bidirectional modulation of anxiety-related and social behaviors by amygdala projections to the medial prefrontal cortex. Neuroscience, 321, 197–209. 10.1016/j.neuroscience.2015.07.041

Felix-Ortiz, A. C., & Tye, K. M. (2014). Amygdala Inputs to the Ventral Hippocampus Bidirectionally Modulate Social Behavior. Journal of Neuroscience, 34(2), 586–595. 10.1523/Jneurosci.4257-13.2014

Fenno, L. E., Mattis, J., Ramakrishnan, C., Hyun, M., Lee, S. Y., He, M., Tucciarone, J., Selimbeyoglu, A., Berndt, A., Grosenick, L., Zalocusky, K. A., Bernstein, H., Swanson, H., Perry, C., Diester, I., Boyce, F. M., Bass, C. E., Neve, R., Huang, Z. J., & Deisseroth, K. (2014). Targeting cells with single vectors using multiple-feature Boolean logic. Nat Methods, 11(7), 763–772. 10.1038/nmeth.2996

Fisher, S. D., Ferguson, L. A., Bertran-Gonzalez, J., & Balleine, B. W. (2020). Amygdala-Cortical Control of Striatal Plasticity Drives the Acquisition of Goal-Directed Action. Curr Biol, 30(22), 4541–4546 e4545. 10.1016/j.cub.2020.08.090

Fremeau, R. T., Jr., Troyer, M. D., Pahner, I., Nygaard, G. O., Tran, C. H., Reimer, R. J., Bellocchio, E. E., Fortin, D., Storm-Mathisen, J., & Edwards, R. H. (2001). The expression of vesicular glutamate transporters defines two classes of excitatory synapse. Neuron, 31(2), 247–260. 10.1016/s0896-6273(01)00344-0

Hintiryan, H., Bowman, I., Johnson, D. L., Korobkova, L., Zhu, M., Khanjani, N., Gou, L., Gao, L., Yamashita, S., Bienkowski, M. S., Garcia, L., Foster, N. N., Benavidez, N. L., Song, M. Y., Lo, D., Cotter, K. R., Becerra, M., Aquino, S., Cao, C., … Dong, H. W. (2021). Connectivity characterization of the mouse basolateral amygdalar complex. Nat Commun, 12(1), 2859. 10.1038/s41467-021-22915-5

Hintiryan, H., Foster, N. N., Bowman, I., Bay, M., Song, M. Y., Gou, L., Yamashita, S., Bienkowski, M. S., Zingg, B., Zhu, M., Yang, X. W., Shih, J. C., Toga, A. W., & Dong, H. W. (2016). The mouse cortico-striatal projectome. Nat Neurosci, 19(8), 1100–1114. 10.1038/nn.4332

Janak, P. H., & Tye, K. M. (2015). From circuits to behaviour in the amygdala. Nature, 517(7534), 284–292. 10.1038/nature14188

Keller, G. B., & Mrsic-Flogel, T. D. (2018). Predictive Processing: A Canonical Cortical Computation. Neuron, 100(2), 424–435. 10.1016/j.neuron.2018.10.003

Kietzman, H. W., Trinoskey-Rice, G., Blumenthal, S. A., Guo, J. D., & Gourley, S. L. (2022). Social incentivization of instrumental choice in mice requires amygdala-prelimbic cortex-nucleus accumbens connectivity. Nat Commun, 13(1), 4768. 10.1038/s41467-022-32388-9

Kim, J., Pignatelli, M., Xu, S., Itohara, S., & Tonegawa, S. (2016). Antagonistic negative and positive neurons of the basolateral amygdala. Nat Neurosci, 19(12), 1636–1646. 10.1038/nn.4414

Kim, J., Zhang, X., Muralidhar, S., LeBlanc, S. A., & Tonegawa, S. (2017). Basolateral to Central Amygdala Neural Circuits for Appetitive Behaviors. Neuron, 93(6), 1464–1479 e1465. 10.1016/j.neuron.2017.02.034

Klavir, O., Prigge, M., Sarel, A., Paz, R., & Yizhar, O. (2017). Manipulating fear associations via optogenetic modulation of amygdala inputs to prefrontal cortex. Nat Neurosci, 20(6), 836–844. 10.1038/nn.4523

Lai, C. W., Shih, C. W., & Chang, C. H. (2022). Analysis of collateral projections from the lateral orbitofrontal cortex to nucleus accumbens and basolateral amygdala in rats. J Neurophysiol, 127(6), 1535–1546. 10.1152/jn.00127.2022

LeDoux, J. (2007). The amygdala. Curr Biol, 17(20), R868–874. 10.1016/j.cub.2007.08.005

Lee, K. M., Ferreira-Santos, F., & Satpute, A. B. (2021). Predictive processing models and affective neuroscience. Neurosci Biobehav Rev, 131, 211–228. 10.1016/j.neubiorev.2021.09.009

Liu, W. Z., Zhang, W. H., Zheng, Z. H., Zou, J. X., Liu, X. X., Huang, S. H., You, W. J., He, Y., Zhang, J. Y., Wang, X. D., & Pan, B. X. (2020). Identification of a prefrontal cortex-to-amygdala pathway for chronic stress-induced anxiety. Nat Commun, 11(1), 2221. 10.1038/s41467-020-15920-7

Lowery-Gionta, E. G., Crowley, N. A., Bukalo, O., Silverstein, S., Holmes, A., & Kash, T. L. (2018). Chronic stress dysregulates amygdalar output to the prefrontal cortex. Neuropharmacology, 139, 68–75. 10.1016/j.neuropharm.2018.06.032

Maccaferri, G., & McBain, C. J. (1996). The hyperpolarization-activated current (Ih) and its contribution to pacemaker activity in rat CA1 hippocampal stratum oriens-alveus interneurones. J Physiol, 497 ( Pt 1)(Pt 1), 119–130. 10.1113/jphysiol.1996.sp021754

Magee, J. C. (1998). Dendritic hyperpolarization-activated currents modify the integrative properties of hippocampal CA1 pyramidal neurons. J Neurosci, 18(19), 7613–7624. 10.1523/JNEUROSCI.18-19-07613.1998

Manoocheri, K., & Carter, A. G. (2022). Rostral and caudal basolateral amygdala engage distinct circuits in the prelimbic and infralimbic prefrontal cortex. Elife, 11. 10.7554/eLife.82688

Massi, L., Hagihara, K. M., Courtin, J., Hinz, J., Muller, C., Fustinana, M. S., Xu, C., Karalis, N., & Luthi, A. (2023). Disynaptic specificity of serial information flow for conditioned fear. Sci Adv, 9(3), eabq1637. 10.1126/sciadv.abq1637

Mate, Z., Poles, M. Z., Szabo, G., Bagyanszki, M., Talapka, P., Fekete, E., & Bodi, N. (2013). Spatiotemporal expression pattern of DsRedT3/CCK gene construct during postnatal development of myenteric plexus in transgenic mice. Cell Tissue Res, 352(2), 199–206. 10.1007/s00441-013-1552-7

Matyas, F., Komlosi, G., Babiczky, A., Kocsis, K., Bartho, P., Barsy, B., David, C., Kanti, V., Porrero, C., Magyar, A., Szucs, I., Clasca, F., & Acsady, L. (2018). A highly collateralized thalamic cell type with arousal-predicting activity serves as a key hub for graded state transitions in the forebrain. Nat Neurosci, 21(11), 1551–1562. 10.1038/s41593-018-0251-9

McDonald, A. J. (1982). Neurons of the lateral and basolateral amygdaloid nuclei: a Golgi study in the rat. J Comp Neurol, 212(3), 293–312. 10.1002/cne.902120307

McDonald, A. J. (1984). Neuronal organization of the lateral and basolateral amygdaloid nuclei in the rat. J Comp Neurol, 222(4), 589–606. 10.1002/cne.902220410

McDonald, A. J. (1992). Projection neurons of the basolateral amygdala: a correlative Golgi and retrograde tract tracing study. Brain Res Bull, 28(2), 179–185. 10.1016/0361-9230(92)90177-y

McDonald, A. J. (1998). Cortical pathways to the mammalian amygdala. Prog Neurobiol, 55(3), 257–332. 10.1016/s0301-0082(98)00003-3

McDonald, A. J., & Culberson, J. L. (1981). Neurons of the basolateral amygdala: a Golgi study in the opossum (Didelphis virginiana). Am J Anat, 162(4), 327–342. 10.1002/aja.1001620404

McGarry, L. M., & Carter, A. G. (2017). Prefrontal Cortex Drives Distinct Projection Neurons in the Basolateral Amygdala. Cell Rep, 21(6), 1426–1433. 10.1016/j.celrep.2017.10.046

Nusser, Z., Naylor, D., & Mody, I. (2001). Synapse-specific contribution of the variation of transmitter concentration to the decay of inhibitory postsynaptic currents. Biophys J, 80(3), 1251–1261. 10.1016/S0006-3495(01)76101-2

O’Leary, T. P., Sullivan, K. E., Wang, L., Clements, J., Lemire, A. L., & Cembrowski, M. S. (2020). Extensive and spatially variable within-cell-type heterogeneity across the basolateral amygdala. Elife, 9. 10.7554/eLife.59003

Paxinos, G., & Franklin, K. B. J. (2004). The Mouse Brain in Stereotaxic Coordinates: Compact Second Edition. Elsevier Science. https://books.google.hu/books?id=EHy1QN1xv0gC

Petrovich, G. D., Canteras, N. S., & Swanson, L. W. (2001). Combinatorial amygdalar inputs to hippocampal domains and hypothalamic behavior systems. Brain Res Brain Res Rev, 38(1-2), 247–289. 10.1016/s0165-0173(01)00080-7

Pinault, D. (1996). A novel single-cell staining procedure performed in vivo under electrophysiological control: morpho-functional features of juxtacellularly labeled thalamic cells and other central neurons with biocytin or Neurobiotin. J Neurosci Methods, 65(2), 113–136. 10.1016/0165-0270(95)00144-1

Pitkanen, A. (2000). Connectivity of the rat amygdaloid complex. In J. P. Aggleton (Ed.), The Amygdala A functional analysis (2nd ed., pp. 31–99). Oxford University Press.

Pitkanen, A., & Amaral, D. G. (1998). Organization of the intrinsic connections of the monkey amygdaloid complex: projections originating in the lateral nucleus. J Comp Neurol, 398(3), 431–458. 10.1002/(sici)1096-9861(19980831)398:3<431::aid-cne9>3.0.co;2-0

Pitkanen, A., Savander, M., Nurminen, N., & Ylinen, A. (2003). Intrinsic synaptic circuitry of the amygdala [Research Support, Non-U.S. Gov’t]. Ann N Y Acad Sci, 985, 34–49. http://www.ncbi.nlm.nih.gov/pubmed/12724146

Pitkanen, A., Savander, V., & LeDoux, J. E. (1997). Organization of intra-amygdaloid circuitries in the rat: an emerging framework for understanding functions of the amygdala. Trends Neurosci, 20(11), 517–523. http://www.ncbi.nlm.nih.gov/entrez/query.fcgi?cmd=Retrieve&db=PubMed&dopt=Citation&list_uids=9364666

Pitkanen, A., Stefanacci, L., Farb, C. R., Go, G. G., LeDoux, J. E., & Amaral, D. G. (1995). Intrinsic connections of the rat amygdaloid complex: projections originating in the lateral nucleus. J Comp Neurol, 356(2), 288–310. 10.1002/cne.903560211

Puelles, L., Kuwana, E., Puelles, E., Bulfone, A., Shimamura, K., Keleher, J., Smiga, S., & Rubenstein, J. L. (2000). Pallial and subpallial derivatives in the embryonic chick and mouse telencephalon, traced by the expression of the genes Dlx-2, Emx-1, Nkx-2.1, Pax-6, and Tbr-1. J Comp Neurol, 424(3), 409–438. 10.1002/1096-9861(20000828)424:3<409::aid-cne3>3.0.co;2-7

Rainnie, D. G., Asprodini, E. K., & Shinnickgallagher, P. (1993). Intracellular-Recordings from Morphologically Identified Neurons of the Basolateral Amygdala. Journal of Neurophysiology, 69(4), 1350–1362. DOI10.1152/jn.1993.69.4.1350

Reppucci, C. J., & Petrovich, G. D. (2016). Organization of connections between the amygdala, medial prefrontal cortex, and lateral hypothalamus: a single and double retrograde tracing study in rats. Brain Struct Funct, 221(6), 2937–2962. 10.1007/s00429-015-1081-0

Rovira-Esteban, L., Peterfi, Z., Vikor, A., Mate, Z., Szabo, G., & Hajos, N. (2017). Morphological and physiological properties of CCK/CB1R-expressing interneurons in the basal amygdala. Brain Struct Funct, 222(8), 3543–3565. 10.1007/s00429-017-1417-z

Sah, P., Faber, E. S., Lopez De Armentia, M., & Power, J. (2003). The amygdaloid complex: anatomy and physiology [Research Support, Non-U.S. Gov’t Review]. Physiol Rev, 83(3), 803–834. 10.1152/physrev.00002.2003

Savander, V., Go, C. G., LeDoux, J. E., & Pitkanen, A. (1995). Intrinsic connections of the rat amygdaloid complex: projections originating in the basal nucleus. J Comp Neurol, 361(2), 345–368. 10.1002/cne.903610211

Scheggia, D., La Greca, F., Maltese, F., Chiacchierini, G., Italia, M., Molent, C., Bernardi, F., Coccia, G., Carrano, N., Zianni, E., Gardoni, F., Di Luca, M., & Papaleo, F. (2022). Reciprocal cortico-amygdala connections regulate prosocial and selfish choices in mice. Nat Neurosci, 25(11), 1505–1518. 10.1038/s41593-022-01179-2

Senn, V., Wolff, S. B., Herry, C., Grenier, F., Ehrlich, I., Grundemann, J., Fadok, J. P., Muller, C., Letzkus, J. J., & Luthi, A. (2014). Long-range connectivity defines behavioral specificity of amygdala neurons [Research Support, Non-U.S. Gov’t]. Neuron, 81(2), 428–437. 10.1016/j.neuron.2013.11.006

Shen, C. J., Zheng, D., Li, K. X., Yang, J. M., Pan, H. Q., Yu, X. D., Fu, J. Y., Zhu, Y., Sun, Q. X., Tang, M. Y., Zhang, Y., Sun, P., Xie, Y., Duan, S., Hu, H., & Li, X. M. (2019). Cannabinoid CB(1) receptors in the amygdalar cholecystokinin glutamatergic afferents to nucleus accumbens modulate depressive-like behavior. Nat Med, 25(2), 337–349. 10.1038/s41591-018-0299-9

Shih, C. W., & Chang, C. H. (2023). Anatomical analyses of collateral prefrontal cortex projections to the basolateral amygdala and the nucleus accumbens core in rats. Brain Struct Funct. 10.1007/s00429-023-02722-y

Shinonaga, Y., Takada, M., & Mizuno, N. (1994). Topographic organization of collateral projections from the basolateral amygdaloid nucleus to both the prefrontal cortex and nucleus accumbens in the rat. Neuroscience, 58(2), 389–397. 10.1016/0306-4522(94)90045-0

Sotres-Bayon, F., Sierra-Mercado, D., Pardilla-Delgado, E., & Quirk, G. J. (2012). Gating of fear in prelimbic cortex by hippocampal and amygdala inputs. Neuron, 76(4), 804–812. 10.1016/j.neuron.2012.09.028

Sprouston, N. S. G. H. M. (2016). Dendrites. In N. S. M. H. G. Stuart (Ed.), Principles of dendritic integration (3rd ed., pp. 351–398). Oxford University Press. 10.1093/acprof:oso/9780198745273.003.0012

Swanson, L. W., & Petrovich, G. D. (1998). What is the amygdala? Trends Neurosci, 21(8), 323–331. 10.1016/s0166-2236(98)01265-x

Tanaka, S., Wu, N., Hsaio, C. F., Turman, J., Jr., & Chandler, S. H. (2003). Development of inward rectification and control of membrane excitability in mesencephalic v neurons. J Neurophysiol, 89(3), 1288–1298. 10.1152/jn.00850.2002

Truitt, W. A., Johnson, P. L., Dietrich, A. D., Fitz, S. D., & Shekhar, A. (2009). Anxiety-like behavior is modulated by a discrete subpopulation of interneurons in the basolateral amygdala. Neuroscience, 160(2), 284–294. 10.1016/j.neuroscience.2009.01.083

Turner, B. H., & Herkenham, M. (1991). Thalamoamygdaloid projections in the rat: a test of the amygdala’s role in sensory processing. J Comp Neurol, 313(2), 295–325. 10.1002/cne.903130208

Tye, K. M., Prakash, R., Kim, S. Y., Fenno, L. E., Grosenick, L., Zarabi, H., Thompson, K. R., Gradinaru, V., Ramakrishnan, C., & Deisseroth, K. (2011). Amygdala circuitry mediating reversible and bidirectional control of anxiety. Nature, 471(7338), 358–362. 10.1038/nature09820

Vereczki, V. K., Muller, K., Krizsan, E., Mate, Z., Fekete, Z., Rovira-Esteban, L., Veres, J. M., Erdelyi, F., & Hajos, N. (2021). Total Number and Ratio of GABAergic Neuron Types in the Mouse Lateral and Basal Amygdala. J Neurosci, 41(21), 4575–4595. 10.1523/JNEUROSCI.2700-20.2021

Vereczki, V. K., Veres, J. M., Muller, K., Nagy, G. A., Racz, B., Barsy, B., & Hajos, N. (2016). Synaptic Organization of Perisomatic GABAergic Inputs onto the Principal Cells of the Mouse Basolateral Amygdala. Front Neuroanat, 10, 20. 10.3389/fnana.2016.00020

Veres, J. M., Nagy, G. A., & Hajos, N. (2017). Perisomatic GABAergic synapses of basket cells effectively control principal neuron activity in amygdala networks. Elife, 6. 10.7554/eLife.20721

Yoon, B. J., Lee, I., Lee, E., Han, N.-E., Slavuj, M., Hwang, J., Lee, A., Sun, T., Jeong, Y., Baik, J.-H., Park, J.-Y., Choi, S.-Y., & Kwag, J. (2023). Persistent enhancement of basolateral amygdala-dorsomedial striatum synapses causes obsessive-compulsive disorder-like behaviors in mice. 10.21203/rs.3.rs-3191969/v1

