## Supplementary figures for "Diversity and connectivity of principal neurons in the lateral and basal nuclei of the mouse amygdala"

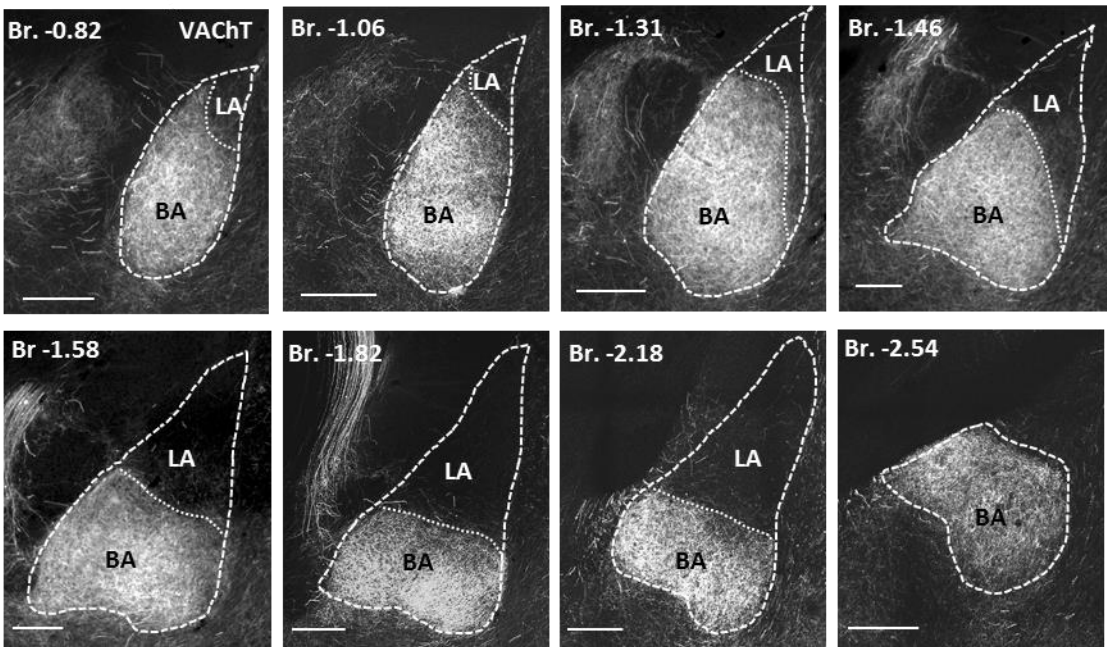


**Suppl. figure 1. Vesicular acetylcholine transporter (VAChT) expression defines the borders between the LA and BA.**

Dashed lines represent the borders of the LA and BA, whereas the dotted lines indicate the borders between the LA and BA. Numbers refer to the distance from Bregma (Br) in mm. Scale bar: 500 µm.

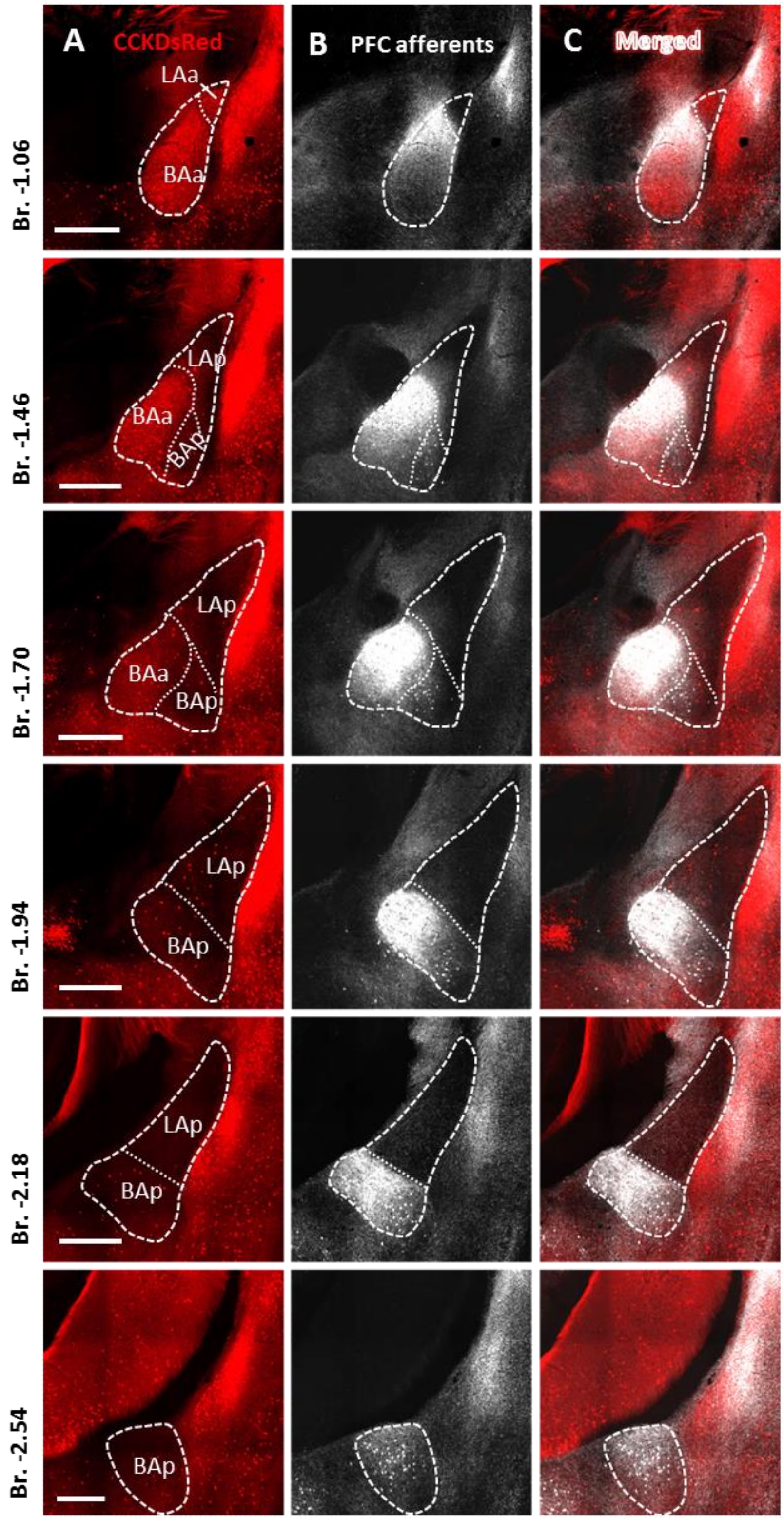


**Suppl. figure 2. Prefrontal cortical afferents in the LA and BA**

Sections containing the basolateral amygdala **(A)** as well as the afferents of prefrontal cortical glutamatergic neurons **(B)** are shown in a VGlut1-Cre::CCK-DsRed double transgenic mouse. Scale bar: 500 um. **(C)** Merged images. Numbers refer to the distance from Bregma (Br) in mm.

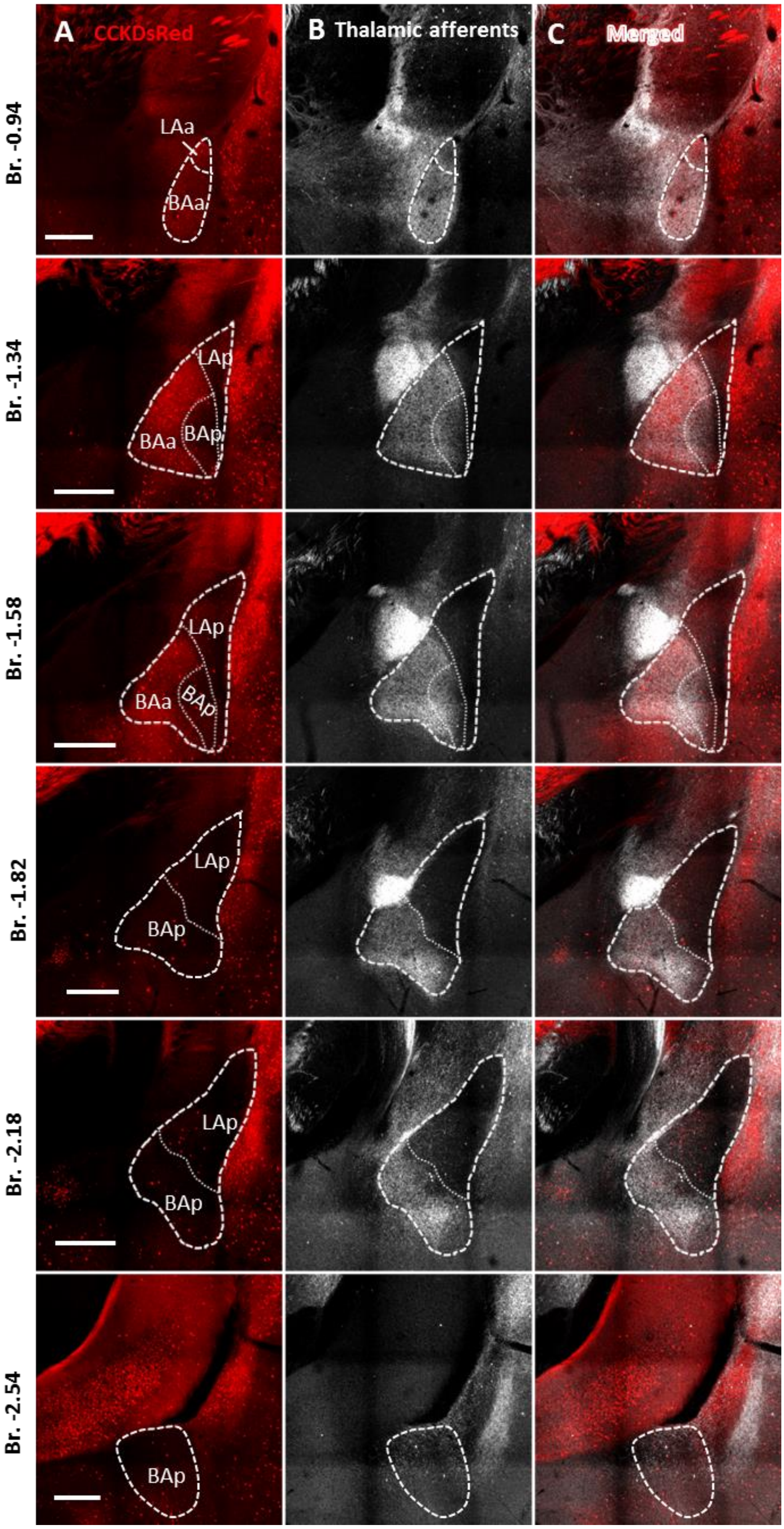
A CCKDsRed
B Thalamic afferents
C Merged
Br. -0.94
Br. -1.34
Br. -1.58
Br. -1.82
Br. -2.18
Br. -2.54
LAa
BAa
LAp
BAp

**Suppl. figure 3. Insular cortical afferents in the LA and BA**

Sections containing the basolateral amygdala **(A)** as well as the afferents of insular cortical glutamatergic neurons **(B)** are shown in a VGlut1-Cre::CCK-DsRed double transgenic mouse. Scale bar: 500 um. **(C)** Merged images. Numbers refer to the distance from Bregma (Br) in mm.

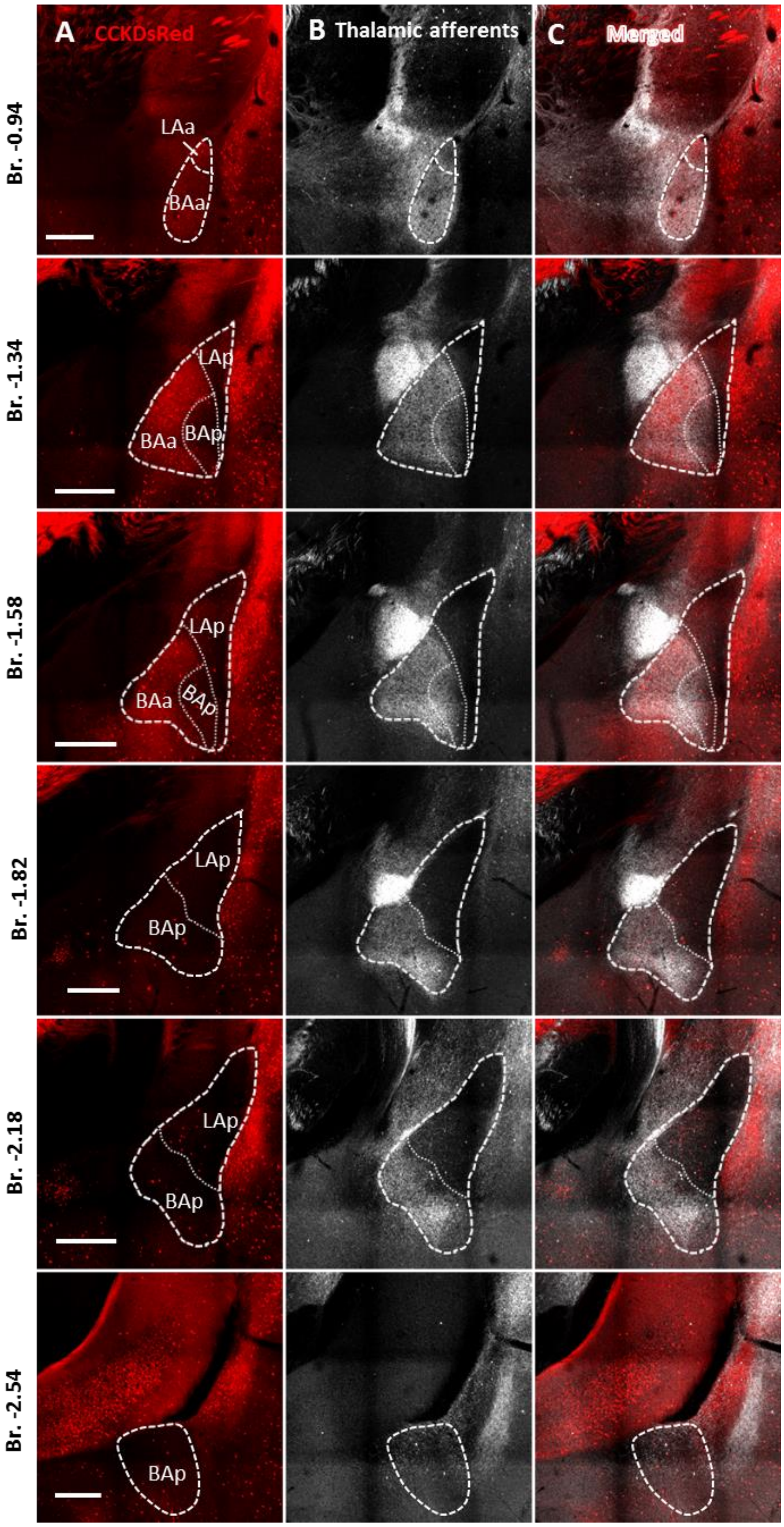


**Suppl. 4. Midline thalamic afferents in the LA and BA**

Sections containing the basolateral amygdala **(A)** as well as the afferents of midline thalamic neurons **(B)** are shown in a CCK-DsRed double transgenic mouse. Scale bar: 500 um. **(C)** Merged images. Numbers refer to the distance from Bregma (Br) in mm.

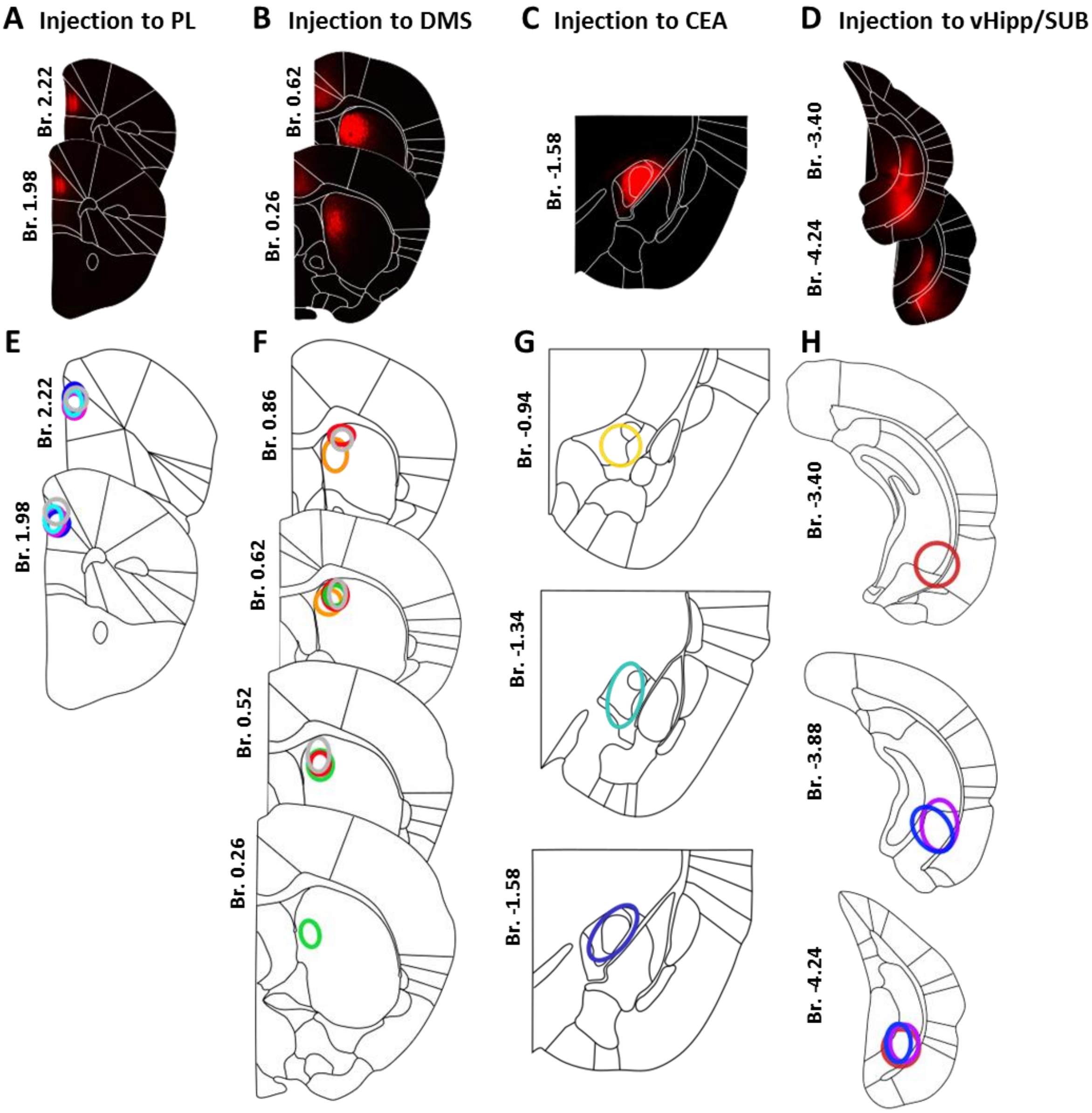


**I**

| Target region | AP | ML | DV | Tracer | Vol. |
|---|---|---|---|---|---|
| **PL** | 1.78/2.1 | 0.3 | 1.0 | AAVrg-Ef1a-IRES-Cre-mCherry | 50-50 nl |
| **DMS** | 0.62/1.1 | 1.3 | 2.3 | AAVrg-Ef1a-IRES-Cre-mCherry or AAVrg-CAG-GFP | 50-50 nl |
| **CEA** | -1.6 | 2.6 | 4.1 | CTB/ | Iontoph. |
| | | | | pAAV-CAG-GFP | 30 nl |
| **vHipp/ SUB** | -3.5/-3.8 | 3.4 | 4.1 | CTB | 25-25 nl |

**Suppl. figure 5. Injection sites of retrograde tracing experiments.**

(**A-D**) Example injection sites targeting the prelimbic cortex (PL), the dorsomedial striatum (DMS), the ventral hippocampus/subiculum (vHipp/SUB) in 2 anteroposterior sections, and the central amygdala (CEA) in 1 anteroposterior section. The atlas drawings were made based on the Allen Brain Atlas (2011). (**E-H**) Location of all retrograde tracer injections used in the study. The core parts of the injections in the same animal are indicated with circles of the same color along different distances from the Bregma (mm). (**I**) Table of stereotaxic coordinates (mm) of the injection sites. The tracers used and the volumes administered are indicated in the table. AP: anteroposterior; Br. Bregma; DV: dorsoventral; ML: mediolateral.

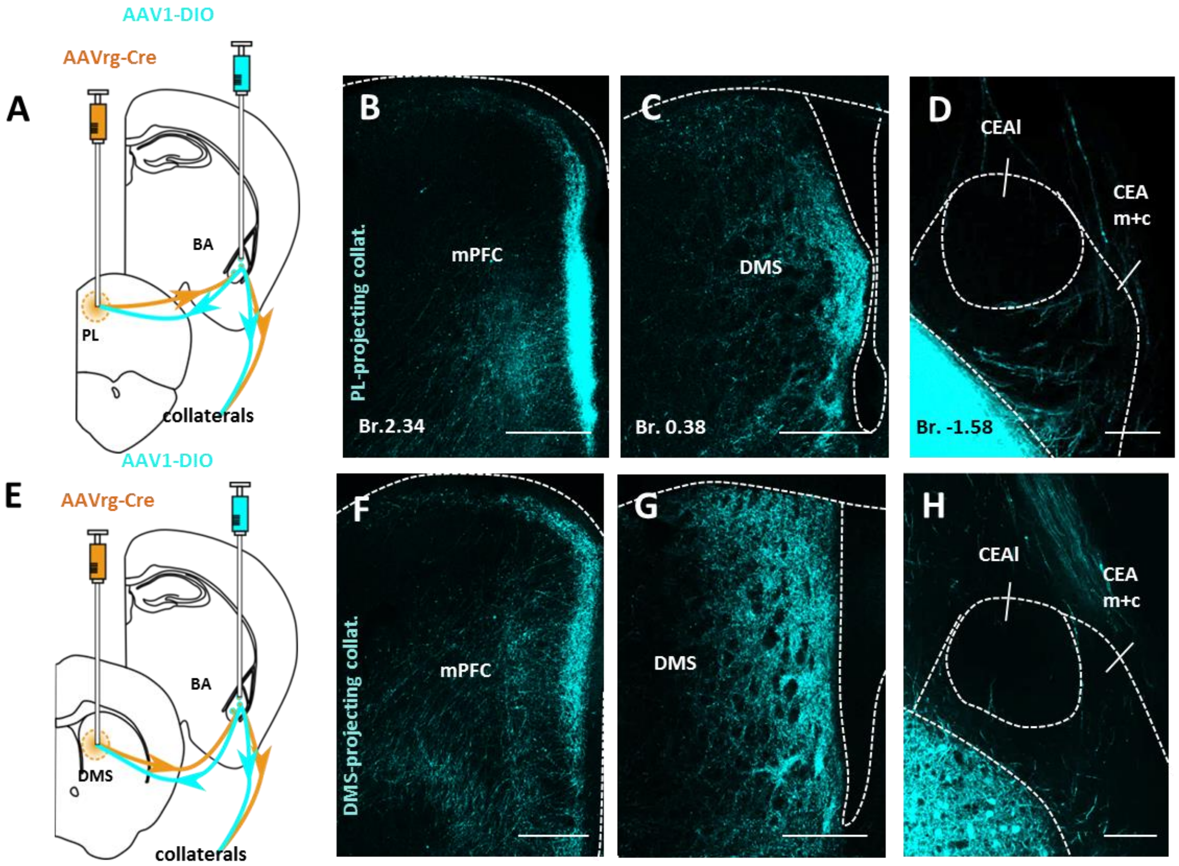


**Suppl. figure 6. Axonal collaterals of DMS- and PL-projecting principal neurons in the mPFC, DMS and CEA**

**(A)** Experimental setup: labeling the axonal collaterals of PL-projecting principal neurons (PNs) using an intersectional approach. **(B-D)** Axonal collaterals of PL-projecting PNs in the mPFC, DMS and CEA. **(E)** Experimental setup: labeling the axonal collaterals of DMS-projecting PNs using an intersectional approach. **(G-H)** Axonal collaterals of DMS-projecting PNs in the mPFC, DMS and CEA. Note that the labeled axons avoid the lateral nucleus of the CEA (CEAl) in both cases. Scale bar: **(B-C, F-G)** 500 µm or **(D, H)** 100 µm. CEAl: central amygdalar nucleus, lateral part; CEAm+c: central amygdalar nucleus, medial and capsular parts; mPFC: medial prefrontal cortex.

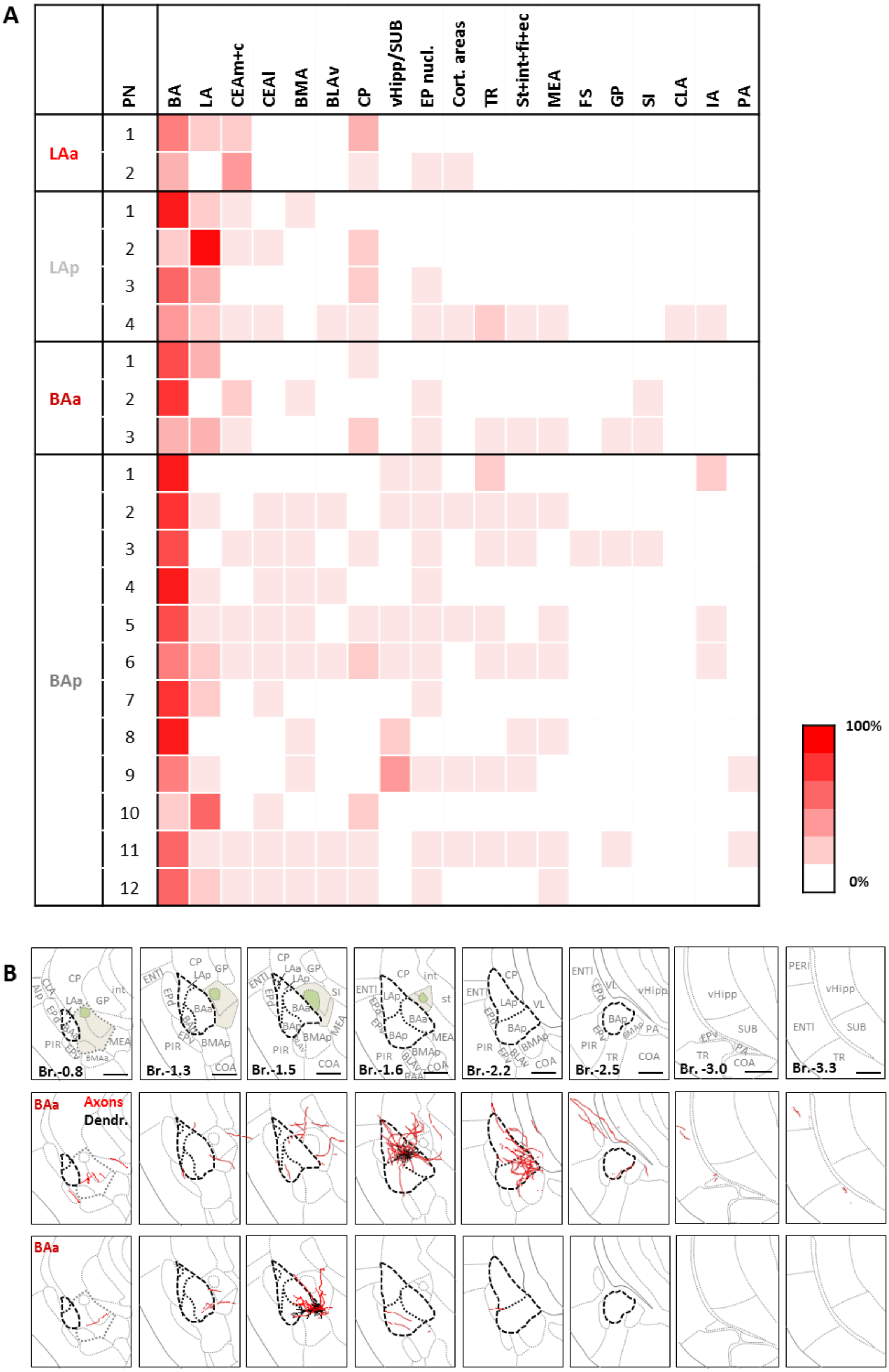
A
PN
BA
LA
CEAm+c
CEAl
BMA
BLAv
CP
vHipp/SUB
EP nucl.
Cort. areas
TR
St+int+fi+ec
MEA
FS
GP
SI
CLA
IA
PA
LAa
LAp
BAa
BAp
1
2
3
4
5
6
7
8
9
10
11
12
100%
0%
B
Br.-0.8
Br.-1.3
Br.-1.5
Br.-1.6
Br.-2.2
Br.-2.5
Br. -3.0
Br.-3.3
CP
GP
int
LAa
LAp
BAa
BAp
BMAp
BMAa
MEA
PIR
EPd
EPv
ENTl
SI
st
VL
COA
TR
PA
vHipp
SUB
PERI
BAa
Axons
Dendr.

**Suppl. figure 7. The presence of axonal collaterals of *in vivo* filled PNs in different amygdalar subnuclei and their surrounding areas**

**(A)** Heatmap of the presence of axonal collaterals of the *in vivo* filled PNs in different regions. **(B)** Two example PNs with soma location in the BAa innervating the medial and capsular subnuclei of the central amygdala (CEA, shown in beige), but avoiding its lateral subnucleus (shown in green). See the List of Abbreviations for the identification of brain areas.

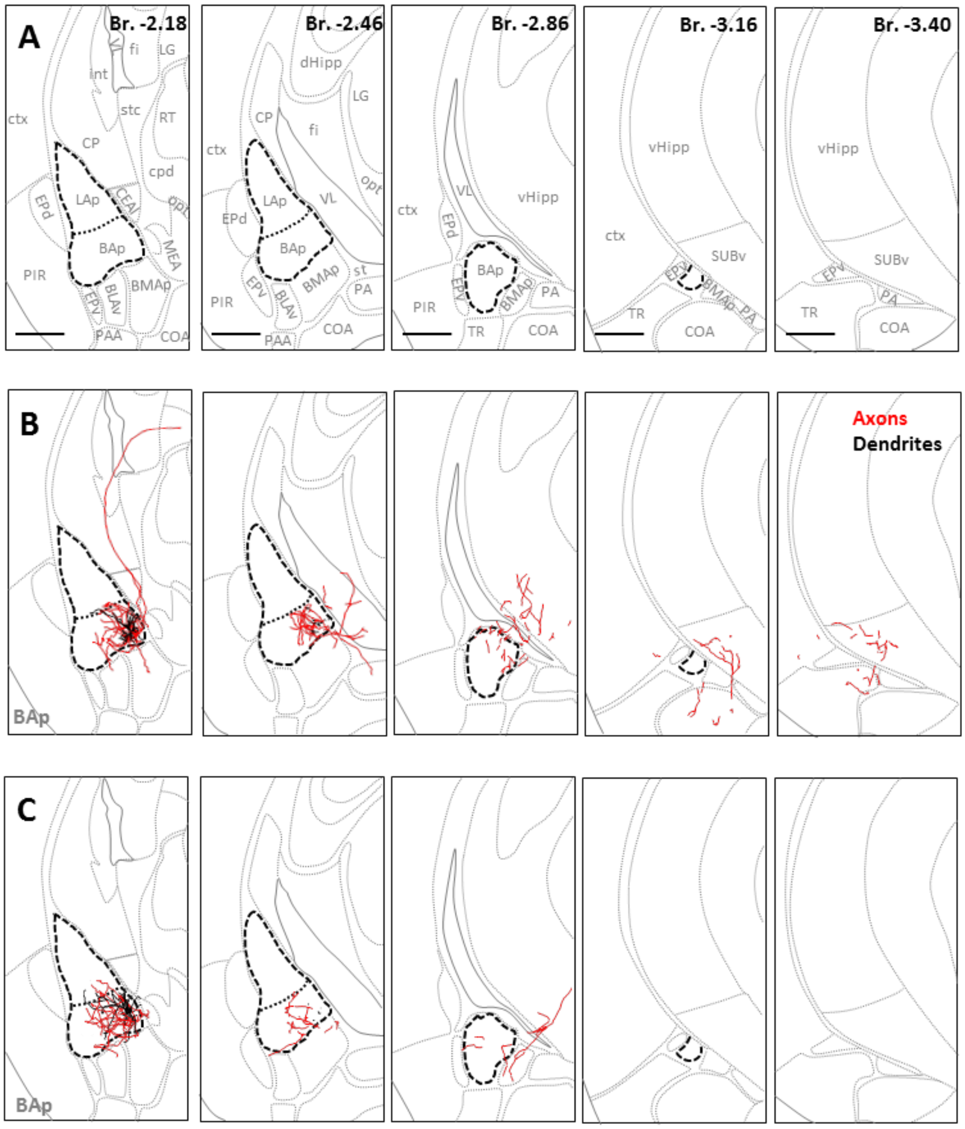


**Suppl. figure 8. *In vivo* filled principal neurons in the medioposterior BA projecting to the different parts of the hippocampus.**

**(A)** Maps showing the LA and BA and the surrounding brain areas along 5 anteroposterior sections based on Allen Brain Mouse Brain Atlas (2011). **(B)** An example principal neuron in the BAp projecting to the dorsal and ventral hippocampus. **(C)** An example principal neuron in the BAp projecting to the ventral hippocampus. Axons are labeled in red and dendrites in black. Scale bar: 500 µm. See the List of Abbreviations for the identification of brain areas.
